# Single-trial Endpoint-summary Measures do not Capture P300 Coupling in the Visual Oddball Paradigm: a Pseudotrial-controlled, Cross-validated Study

**DOI:** 10.64898/2025.12.17.694588

**Authors:** Erkan Biber

**Author notes:** **Corresponding Author:** Erkan Biber, Institute of Biomedical Engineering, Boğaziçi University, Kandilli Campus, 34684 Çengelköy, Istanbul, Turkey.

## Abstract

Single-trial analysis of event-related potentials promises access to the trial-to-trial variability that averaging discards, and many studies report early-window summary measures that covary with later component amplitudes. Such couplings can, however, arise from the temporal autocorrelation of continuous EEG rather than from stimulus-locked processing. We asked whether the conventional family of endpoint-summary measures those that collapse a time window to a single value, including mean amplitude, root-mean-square, variance, signal-complexity measures (permutation entropy, sample entropy, Lempel-Ziv complexity), and Hjorth parameters, captures genuine stimulus-locked information about P300 amplitude in the active visual oddball once autocorrelation is controlled. Analyzing the ERP CORE visual P3 dataset (*N* = 27; 1,084 trials, 213 target and 871 standard, with experimental condition as a covariate), we related each early-window (0-150 ms) measure to P300 amplitude at Pz and re-estimated every model on pseudotrials placed at random latencies in the same recording; the direction of change under this substitution, not the raw effect size, is the diagnostic. Cross-channel amplitude and energy couplings strengthened under pseudotrial substitution, indicating dependence on background structure. Large same-channel coupling (*R*² ≈ 0.31) was unchanged under substitution and present at every electrode, including the eye channels, identifying it as general within-trial temporal continuity rather than a P300-specific process. Complexity measures carried near-zero population-level coupling but large, directionally split per-subject slopes. An independent dataset (different laboratory and hardware; same paradigm) reproduced the same-channel continuity result (*N* = 90 participants; 83 for pseudotrial fits) and the directionally split per-subject pattern across complexity measures. Endpoint-summary measures therefore do not capture consistent population-level P300 coupling once autocorrelation is controlled; the complexity family carries person-specific coupling that cancels at the population level, motivating analytic designs sensitive to individual differences.

## 1. Introduction

### 1.1 Trial-to-trial variability in event-related potentials

Event-related potentials (ERPs) are scalp-recorded voltage changes time-locked to sensory, cognitive, or motor events, and they have been a central tool of cognitive neuroscience for more than half a century (Luck, 2014). Their standard derivation rests on a simple premise: average many repeated trials so that activity unrelated to the event cancels, leaving a stable waveform whose peaks and troughs can be measured and compared. The procedure is powerful, but it is also lossy. Whatever varies from one trial to the next, fluctuations in arousal, attention, expectation, or the moment-to-moment state of the underlying networks, is treated as noise to be averaged away. For much of the history of the field this was a reasonable price to pay, because the averaged waveform was the object of interest. Over the past two decades, however, attention has increasingly turned to the discarded variability itself, on the hypothesis that the trial-to-trial modulation of an ERP carries information about the cognitive and neural state of the participant on that particular trial (Pernet et al., 2011; Stokes & Spaak, 2016).

The P300 (or P3) has been a natural focus of this effort. Elicited reliably by infrequent, task-relevant stimuli in the oddball paradigm, it is among the largest and most robust components in the human ERP, and its amplitude is sensitive to attention, stimulus probability, and the updating of working-memory representations (Polich, 2007). Its size makes it tractable at the single-trial level in a way that smaller components are not, and its functional sensitivity makes single-trial variation in its amplitude scientifically interesting: if the P300 indexes the allocation of attentional or memory resources, then trial-by-trial fluctuation in its amplitude might track corresponding fluctuation in those processes.

### 1.2 Early-window summary measures and the promise of single-trial prediction

A recurring strategy in this literature is to summarize the early portion of a single trial, the pre-component window, before the P300 itself has developed, with a scalar feature, and to ask whether that feature covaries with the amplitude of the later component on the same trial. The intuition is that the state of the system in the early window, captured by some property of the signal, shapes or forecasts the response that follows. A wide range of such features has been proposed. Some are drawn from the amplitude domain: the mean voltage, the root-mean-square, the variance of the early window. Others are intended to capture the *shape* or *complexity* of the early signal rather than its magnitude, measures such as permutation entropy, sample entropy, and Lempel–Ziv complexity, which quantify the regularity or predictability of a time series and have been applied to EEG as putative indices of neural state (Bachiller et al., 2015; Bruña et al., 2012). What these features share is that each collapses a time window into a single number. We refer to them collectively as *endpoint-summary measures*: instruments that reduce the early-window trajectory to one value and relate that value to a later endpoint.

The appeal of this approach is its simplicity and its compatibility with standard regression machinery. A single early-window scalar per trial, a single P300 amplitude per trial, and a mixed-effects model relating the two across trials and participants, the analysis is straightforward to implement and to interpret. Recovering a component’s amplitude on individual trials is itself a well-developed enterprise, with a substantial methodological literature on single-trial estimation and classification of ERPs (Blankertz, Lemm, Treder, Haufe, & Müller, 2011), and single-trial P300 amplitude has been shown to covary with trial-level variables such as reaction time and the preceding stimulus sequence (Holm, Ranta-aho, Sallinen, Karjalainen, & Müller, 2006). Against this backdrop the literature contains many reports of early-window couplings, framed as evidence that the early window predicts, encodes, or carries information about the component that follows.

### 1.3 The autocorrelation problem

There is a structural reason to treat these reports with caution. Continuous EEG is strongly autocorrelated: the voltage at one moment is highly predictable from the voltage shortly before, both because of the low-pass characteristics of volume conduction and filtering and because of the broadband 1/f structure of the background signal (He, 2014; Voytek et al., 2015). Any two windows extracted from the same ongoing recording will therefore tend to be correlated simply because they are drawn from the same temporally continuous process, with no need for any stimulus-locked mechanism linking them. When an early-window summary and a later component amplitude are both measured from the same epoch of the same electrode, a regression relating them can be positive and statistically reliable even if nothing about the stimulus or the cognitive response is involved, the relationship can be a property of the signal’s autocorrelation rather than of neural processing. This concern is sharpened by recent evidence that the P300 itself may arise, at least in part, not as a discrete additive deflection but through a baseline-shift mechanism in which the stimulus-driven modulation of ongoing alpha oscillations with a non-zero mean produces the apparent positivity (Studenova et al., 2023). To the extent that this is so, the later “component” and the earlier window are expressions of the same continuously evolving oscillatory signal, and a within-electrode coupling between them is exactly what background structure would predict.

This is not a hypothetical concern. It has direct analogues in adjacent literatures. In the study of heartbeat-evoked potentials, apparent relationships between pre-stimulus cardiac-locked activity and task-evoked responses have been shown to be substantially or wholly attributable to background activity once appropriate controls are applied (Steinfath et al., 2025). The methodological lesson generalizes: when a coupling is measured within a single continuous recording, the burden is on the analyst to show that the coupling reflects the event of interest rather than the temporal structure that would be present regardless.

### 1.4 Pseudotrial control

A direct way to discharge that burden is the pseudotrial control (Steinfath et al., 2025). The logic is to construct surrogate “trials” that share the background temporal structure of the real data but are not time-locked to any stimulus, windows placed at random latencies in the continuous recording, away from genuine events, and then subjected to the identical processing and measurement pipeline. The same coupling analysis is run on these pseudotrials. Because the pseudotrials retain the autocorrelation structure of the signal but lack any stimulus-locked component, the comparison is diagnostic. If a coupling is driven by the signal’s background structure, it will be preserved or even strengthened when real trials are replaced by pseudotrials. If a coupling reflects genuine stimulus-locked processing, it should weaken under the substitution, because the stimulus-locked component that produced it is absent from the surrogates. The *direction of change* in the coupling under pseudotrial substitution, rather than its raw magnitude, is therefore the informative quantity.

### 1.5 The present study

We apply this logic systematically to the endpoint-summary family in the active visual oddball paradigm. Using the ERP CORE visual P3 dataset (Kappenman et al., 2021) as a primary sample and an independent dataset collected with the same paradigm at a different laboratory (Isbell et al., 2025) as a cross-validation sample, we ask three questions. First, for each endpoint-summary measure relating an early window to the later P300, is the coupling preserved or weakened under pseudotrial substitution, that is, is it a signature of background autocorrelation or of stimulus-locked processing? Second, where a coupling survives, is it specific to the parietal sites where the P300 is generated, or is it a general property of any single electrode? Third, for measures whose population-level coupling is weak or absent, does that population-level result conceal systematic individual-level structure that an average obscures? The answers, taken together, bear directly on whether the endpoint-summary family is an adequate instrument for single-trial characterization of the P300, and on what analytic designs are required to make progress.

## 2. Methods

### 2.1 Design overview

The analysis related early-window single-trial summary measures to later P300 amplitude, with every coupling estimate accompanied by a pseudotrial control estimate. The same pipeline was applied to a primary dataset and an independent cross-validation dataset. Three families of measure were examined: amplitude/position/energy measures, a same-channel temporal-continuity model, and signal-complexity measures. For the complexity family, a per-subject individual-differences analysis was added to the population-level mixed-effects analysis.

### 2.2 Primary dataset

The primary dataset was the visual P3 component of the ERP CORE resource (Kappenman et al., 2021), an openly available compendium of optimized paradigms and recordings for seven standard ERP components. The visual P3 task is an active visual oddball in which participants view a stream of the letters A–E and respond according to which letter is the designated target on a given block, so that each letter serves as target on some blocks and as standard on others. Recordings were obtained from 27 of the 40 ERP CORE participants who met the trial-count criterion described below. Both target and standard trials were retained, yielding 1,084 epochs (213 target and 871 standard, the ∼1:4 ratio of the oddball design) across the retained participants (approximately 40 per participant); experimental condition was entered as a covariate in every coupling model, so that the analyses concern trial-to-trial covariation of the P300 rather than the target–standard difference.

### 2.3 Cross-validation dataset

The cross-validation dataset was OpenNeuro ds006018 (Isbell et al., 2025), an EEG dataset collected from 127 young adults using tasks directly acquired or adapted from ERP CORE, including the visual P3 oddball with the same A–E letter design.

It provides a stringent test of whether any coupling identified in the primary sample is a property of the paradigm and the signal rather than of one laboratory’s data. 90 participants met the trial-count criterion and were retained for analysis (3,130 target trials for the real-trial models); the matched-pseudotrial fits used for the pseudotrial control retained 83 of these participants (2,270 pseudotrials), the small reduction reflecting participants for whom too few clean pseudotrials could be placed within the continuous recording. The same 90 participants entered the per-subject individual-differences analysis. The dataset is distributed under a CC0 licence and was accessed programmatically via EEGDash from the OpenNeuro data store.

The two datasets share the visual P3 paradigm but differ in population, recording hardware, and montage, which is what makes the cross-validation informative; their characteristics are compared in Table 1.

**Table 1.** Characteristics of the primary and cross-validation datasets. Both use the ERP CORE active visual oddball (letters A–E); the differences in recording system, montage, and population are what make the cross-validation a stringent test of generalization.

| Characteristic | Primary: ERP CORE | Cross-validation: ds006018 |
| --- | --- | --- |
| Source | Kappenman et al. (2021) | Isbell et al. (2025) |
| Paradigm | Active visual oddball (letters A–E) | Active visual oddball (letters A–E) |
| Total participants | 40 | 127 |
| Retained (trial-count criterion) | 27 | 90 / 83 (matched-pseudotrial fits) |
| Population | Neurotypical young adults | Socioeconomically diverse young adults, |
| Age range | 18–30 | 18–30 |
| Recording system | Biosemi ActiveTwo (active electrodes) | Brain Products actiCHamp Plus (active electrodes) |
| Scalp electrodes | 30 | 26 (32-channel cap) |
| Sampling rate | 1024 Hz, 24-bit | 500 Hz |
| Early-window samples (0–150 ms) | ≈ 154 | 75 |
| P300 electrode | Pz | Pz |
| Trials analysed | Target + standard (213 + 871; condition covariate) | Target only |

### 2.4 Preprocessing

Both datasets were processed with an identical pipeline implemented in MNE-Python (Gramfort et al., 2013). Continuous EEG was band-pass filtered between 0.1 and 30 Hz with a zero-phase FIR filter (firwin design, Hamming window; Widmann et al., 2015) and re-referenced to the average of the scalp electrodes, excluding the electrooculogram (EOG) channels.

Ocular artifacts were removed by independent component analysis (FastICA; Hyvärinen & Oja, 2000). Following best practice for ICA stability (Winkler et al., 2015), the decomposition was fitted on a copy of the continuous data high-pass filtered at 1 Hz, retaining components that explained 99% of the variance, and components correlated with the electrooculogram (EOG) channels were identified automatically by MNE’s *find_bads_eog* routine and removed from the data filtered at 0.1 Hz.

In the cross-validation dataset, the horizontal- and vertical-eye channels were assigned the EOG channel type and the mastoid channels were designated non-scalp before ICA, so that the decomposition operated only on scalp activity. Where no ocular component met the correlation criterion, no component was removed for that participant.

Continuous data were then segmented into stimulus-locked epochs and baseline-corrected. Epochs were subjected to peak-to-peak artifact rejection with a threshold of ±100 µV applied across all scalp channels; epochs exceeding this threshold on any channel were discarded. The identical rejection criterion was applied to real trials and pseudotrials (Section 2.8). In the primary dataset both target and standard trials were retained, selected by their event codes, with experimental condition entered as a covariate in every coupling model; in the cross-validation dataset, which provides only target events for this paradigm, analysis was restricted to target trials.

Participants retaining fewer than the minimum number of clean trials were excluded, a criterion that produced the *N* = 27 primary sample; in the cross-validation dataset the retained count was 90 for the real-trial and per-subject analyses, with 83 of those participants retained for the matched-pseudotrial fits (Section 2.3). A trial-count diagnostic confirmed the expected composition in each dataset: the ∼1:4 target-to-standard ratio in the primary sample (213 target, 871 standard) and the target-only structure of the cross-validation sample.

### 2.5 Early-window summary measures

For each retained trial, summary measures were computed on the early window, defined as 0–150 ms post-stimulus, before the onset of the P300. Because the two datasets were recorded at different sampling rates (1024 Hz for ERP CORE, 500 Hz for ds006018), this fixed time window corresponded to a different number of samples in each (approximately 154 and 75, respectively); the window was defined in milliseconds rather than samples so that the measured interval was identical across datasets. The amplitude/position/energy family comprised the mean amplitude, median, root-mean-square, and variance of the early window, together with measures extended in robustness analyses (median absolute deviation, peak-to-peak amplitude, skewness, kurtosis, slope, and the Hjorth mobility and complexity parameters). The signal-complexity family comprised permutation entropy (M13; Bandt & Pompe, 2002; embedding dimension 3, time delay 1, normalised), sample entropy (M14; Richman & Moorman, 2000), and Lempel–Ziv complexity (M15; Lempel & Ziv, 1976; computed on the median-thresholded binary sequence of the window, normalized), each computed with the *antropy* library (Vallat, 2023) and quantifying a different aspect of the regularity or predictability of the early-window time series. Permutation entropy characterizes the diversity of ordinal patterns in the signal; sample entropy measures the conditional probability that similar subsequences remain similar as the embedding length increases; and Lempel–Ziv complexity counts the number of distinct substrings in a symbolic recoding of the signal. All measures were standardized before entry into the models.

The three amplitude/energy measures of a window are not independent but are linked by an exact algebraic identity. For any time window, the root-mean-square (RMS), the mean amplitude, and the standard deviation (SD) satisfy

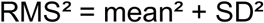

The RMS combines the signed direct-current (mean) level of the window with its mean-centered variability (SD). In the competitive models that enter the mean and SD of the same window as predictors of the P300 (M4a/M4b for the frontal window, M9a/M9b for the parietal window; Section 2.7), the mean and SD covariates are therefore obtained from the RMS formula, they are the two algebraic components into which the RMS of the same window decomposes (mean = signed level, SD = mean-centered energy), rather than independently defined quantities. Entering them jointly partitions the total early-window energy indexed by the RMS into its signed-level and variability parts, so that each competitive coefficient reports the unique contribution of one component with the other held constant.

### 2.6 P300 measurement

The dependent measure was the mean amplitude of the P300 in the 300–600 ms window at the parietal electrode Pz, consistent with the canonical centro-parietal topography of the target-evoked P3b (Polich, 2007). For the same-channel and full-montage analyses described below, the corresponding later-window mean amplitude was measured at each electrode in turn.

### 2.7 Coupling models

Single-trial coupling was estimated with linear mixed-effects models (Baayen, Davidson, & Bates, 2008) fitted by restricted maximum likelihood in statsmodels (Seabold & Perktold, 2010). Before modelling, both the predictor and the outcome were standardized within each participant using a robust z-transformation, each observation centered on the participant’s median and divided by 1.4826 times the participant’s median absolute deviation, which removes between-participant differences in scale and offset while remaining insensitive to outlying trials. Each model related a single standardized early-window measure to the standardized P300 amplitude with a by-participant random intercept and no random slope:

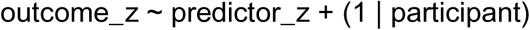

Coupling magnitude is reported as the standardized fixed-effect coefficient (β) and the associated marginal *R*², computed by the variance-decomposition method of Nakagawa and Schielzeth (2013) as the fixed-effect variance divided by the sum of fixed-effect, random-effect, and residual variance. The random-intercept specification was used uniformly across the real-trial and pseudotrial fits so that the two estimates are directly comparable on the same statistical basis. The choice of a random-intercept rather than a random-intercept-plus-random-slope structure was confirmed by model comparison: for every headline model, the random-intercept fit was preferred over the random-slope fit by the Akaike information criterion (ΔAIC in favor of the random-intercept structure of 32–54), and residual diagnostics (quantile–quantile and residuals-versus-fitted plots for representative models across all model families) showed no material departures from the model assumptions (Supplementary Figure S1).

The full set of models is given in Table 2. They fall into four groups. Cross-channel models relate an early-window measure at the frontal electrode Fz to the later P300 at Pz, testing whether early activity at one location forecasts the parietal component. Same-channel models relate an early-window measure at Pz to the later P300 at the same electrode, testing whether the early and late portions of the same parietal signal are coupled. Complexity models relate early-window signal-complexity measures at Fz to the P300. Distributional-shape, robust-amplitude, trend, and Hjorth models extend the same logic across the remaining endpoint-summary measures.

**Table 2.** The full set of single-trial coupling models. Each model relates one early-window measure (at the electrode shown) to the later P300 mean amplitude at Pz (300–600 ms). M1–M12 are the amplitude and energy models; M13–M15 the signal-complexity measures; M16–M23 the distributional-shape, robust-amplitude, trend, and Hjorth extensions. “Competitive” denotes a model in which a second early-window measure is entered as a covariate to isolate the focal predictor.

| Model | Label | Window | Site<br>(early →<br>P3) | Measure | What it indexes | Type |
| --- | --- | --- | --- | --- | --- | --- |
| M1 | Frontal RMS | 0–150<br>ms | Fz → Pz | Root-mean-square | Overall energy (magnitude irrespective of sign) of early frontal activity | Cross-channel |
| M2 | Frontal mean | 0–150<br>ms | Fz → Pz | Mean amplitude | Net signed deflection (DC level) of the early frontal response | Cross-channel |
| M3 | Frontal SD | 0–150<br>ms | Fz → Pz | Standard deviation | Within-trial frontal variability, independent of mean level | Cross-channel |
| M4a | Frontal mean, competitive | 0–150<br>ms | Fz → Pz | Mean (SD covariate) | Unique contribution of mean level, frontal variability held constant | Cross-channel |
| M4b | Frontal SD, competitive | 0–150<br>ms | Fz → Pz | SD (mean covariate) | Unique contribution of variability, mean level held constant | Cross-channel |
| M5 | Frontal RMS | 0–200<br>ms | Fz → Pz | Root-mean-square | M1 at a longer window; window-length dependence of energy coupling | Cross-channel |
| M6 | Frontal RMS | 0–250<br>ms | Fz → Pz | Root-mean-square | M1 at a longer window; window-length dependence of energy coupling | Cross-channel |
| M5m | Frontal mean | 0–200<br>ms | Fz → Pz | Mean amplitude | M2 at a longer window; window-length dependence of mean coupling | Cross-channel |
| M6m | Frontal mean | 0–250<br>ms | Fz → Pz | Mean amplitude | M2 at a longer window; window-length dependence of mean coupling | Cross-channel |
| M7 | Frontal baseline RMS | –150–0<br>ms | Fz → Pz | Root-mean-square | Pre-stimulus energy; cannot reflect evoked activity (autocorrelation reference) | Baseline control |
| M7m | Frontal baseline mean | –150–0<br>ms | Fz → Pz | Mean amplitude | Pre-stimulus level; autocorrelation-background reference | Baseline control |
| M8 | Parietal RMS | 0–150 | Pz → Pz | Root-mean- | Same-channel energy | Same- |

| **Model** | **Label** | **Window**<br>ms | **Site**<br>(early →<br>P3) | **Measure** | **What it indexes** | **Type** |
| --- | --- | --- | --- | --- | --- | --- |
|  |  |  |  | square | coupling at the P3 electrode | channel |
| M9a | Parietal mean, competitive | 0–150 ms | Pz → Pz | Mean (SD covariate) | Largest coupling in the dataset; same-channel mean level, examined in greatest detail | Same-channel |
| M9b | Parietal SD, competitive | 0–150 ms | Pz → Pz | SD (mean covariate) | Companion to M9a; unique contribution of parietal variability | Same-channel |
| M10 | Parietal baseline RMS | –150–0 ms | Pz → Pz | Root-mean-square | Pre-stimulus same-channel energy; autocorrelation reference | Baseline control |
| M11 | Parietal RMS + baseline | 0–150 ms | Pz → Pz | RMS (baseline covariate) | Whether same-channel energy coupling survives adjustment for pre-stimulus level | Same-channel |
| M12 | Parietal mean + baseline | 0–150 ms | Pz → Pz | Mean (baseline covariate) | Whether same-channel mean coupling survives adjustment for pre-stimulus level | Same-channel |
| M13 | Permutation entropy | 0–150 ms | Fz → Pz | Permutation entropy | Diversity of ordinal patterns; temporal complexity of early activity | Complexity |
| M14 | Sample entropy | 0–150 ms | Fz → Pz | Sample entropy | Conditional regularity (likelihood similar subsequences stay similar) | Complexity |
| M15 | Lempel–Ziv complexity | 0–150 ms | Fz → Pz | Lempel–Ziv complexity | Number of distinct substrings in a symbolic recoding; signal compressibility | Complexity |
| M16 | Frontal skewness | 0–150 ms | Fz → Pz | Skewness | Asymmetry of the early-window amplitude distribution | Distributional shape |
| M17 | Frontal kurtosis | 0–150 ms | Fz → Pz | Kurtosis | Peakedness / tail weight of the early-window distribution | Distributional shape |
| M18 | Frontal median | 0–150 ms | Fz → Pz | Median | Robust analogue of M2 (mean level); robustness check | Robust amplitude |
| M19 | Frontal MAD | 0–150 ms | Fz → Pz | Median absolute deviation | Robust analogue of M3 (variability); robustness check | Robust amplitude |

| **Model** | **Label** | **Window** | **Site**<br>**(early →**<br>**P3)** | **Measure** | **What it indexes** | **Type** |
| --- | --- | --- | --- | --- | --- | --- |
| M20 | Frontal peak-to-peak | 0–150 ms | Fz → Pz | Peak-to-peak amplitude | Full range (max – min) of early frontal activity | Robust amplitude |
| M21 | Frontal slope | 0–150 ms | Fz → Pz | Within-window slope | Linear trend (direction of drift); boundary between endpoint and trajectory | Trend |
| M22 | Frontal Hjorth mobility | 0–150 ms | Fz → Pz | Hjorth mobility | Mean frequency of early frontal activity | Hjorth |
| M23 | Frontal Hjorth complexity | 0–150 ms | Fz → Pz | Hjorth complexity | Bandwidth / frequency spread of early frontal activity | Hjorth |

Two model variants recur within these groups: *competitive* models (M4a/M4b, M9a/M9b) include a second early-window measure as a covariate to isolate the focal predictor’s unique contribution, and *baseline-covariate* models (M11, M12) test whether coupling survives adjustment for the pre-stimulus signal level. Baseline-window models (M7, M7m, M10), placed before stimulus onset, cannot reflect evoked activity and therefore serve as a reference for the level of autocorrelation-driven background coupling.

### 2.8 Pseudotrial control

Every model was re-estimated on pseudotrials. For each participant, pseudotrials were generated by placing epoch windows at random latencies within the same continuous recording, matched in number to that participant’s retained real trials, constrained to fall no closer than a minimum gap from any genuine stimulus event, and passed through the identical baseline correction, artifact rejection, and measurement steps applied to real trials. Each model was then re-estimated using these pseudotrials as the data source. The logic is directional: a coupling arising from stimulus-locked neural processing should weaken when stimulus-locked windows are replaced by randomly placed ones, whereas a coupling arising from the signal’s autocorrelation structure should be preserved or strengthened, since that structure is present throughout the continuous recording.

To ensure that the inference did not depend on pseudotrial placement parameters, four configurations were examined, formed by crossing two minimum-gap values (0.5 s and 1.0 s) with two peak-to-peak rejection thresholds (±100 µV and ±150 µV).

These parameters trade off against the number of usable pseudotrials: a longer gap and a stricter threshold each reject more candidate windows, shrinking the surrogate sample and so weakening the basis for comparison with the full real-trial set.

The configuration combining the 0.5 s gap with the ±150 µV threshold (the loosest of the four that still keeps pseudotrials clear of stimulus-locked activity and removes gross artifacts) retained the largest pseudotrial sample, approaching the real-trial count and all 27 participants, against 13–43% of trials and only 10–20 participants for the stricter configurations (Table 3), and was therefore used as the primary comparison because it affords the most statistically powerful test, not because it yields a different result. Across all four configurations the conclusions were unchanged: the same-channel coupling remained large with its coefficient preserved (M9a pseudotrial *R*² 0.23–0.30 against a real-trial 0.31 in every configuration), and the cross-channel coupling remained negligible (*R*² ≤ 0.012 throughout). Pseudotrial placement used a fixed random seed (12345) for reproducibility.

**Table 3.** The four pseudotrial configurations and their behavior. Retention is the pseudotrial trial count as a percentage of the real-trial set (1,084 trials, 27 participants). M9a is the same-channel temporal-continuity model (real-trial *R*² = 0.310); M1 is the frontal cross-channel energy model (real-trial *R*² = 0.007). The same-channel coupling is preserved and the cross-channel coupling negligible under every configuration; Config 4 is the primary comparison because it retains the largest, most representative pseudotrial sample.

| Configuration | Min gap | Threshold | Pseudotrials (% of real) | Participants | M9a $R^2_{\text{pseudo}}$ | M1 $R^2_{\text{pseudo}}$ |
| --- | --- | --- | --- | --- | --- | --- |
| 1 | 1.0 s | $\pm 100 \mu\text{V}$ | 157 (14%) | 11 | 0.298 | 0.004 |
| 2 | 0.5 s | $\pm 100 \mu\text{V}$ | 455 (42%) | 16 | 0.279 | 0.002 |
| 3 | 1.0 s | $\pm 150 \mu\text{V}$ | 426 (39%) | 20 | 0.303 | 0.012 |
| 4 (primary) | 0.5 s | $\pm 150 \mu\text{V}$ | 952 (88%) | 27 | 0.229 | 0.001 |

For each model, the coupling coefficient and *R*² were compared between the real-trial and pseudotrial fits. The primary diagnostic was the **autocorrelation ratio (AUR)**, defined as the pseudotrial-to-real coupling ratio:

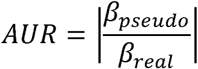

that is, the magnitude of the pseudotrial coupling divided by the magnitude of the real-trial coupling, equivalently can be summarized by the ratio of pseudotrial to real-trial *R*² but in this study beta based AUR is preferred because it preserves the direct linear effect size of the coupling, whereas R^2^-based AUR squares the effect and can exaggerate differences between already small associations (for comparison see Table 8). An AUR at or above 1 indicates a coupling carried by background autocorrelation (the coupling is as strong or stronger when the stimulus-locked response is removed), whereas an AUR well below one indicates a stimulus-locked contribution (the coupling weakens when stimulus locking is removed). The AUR, rather than the raw effect size, is the informative quantity throughout the analyses below, and is reported for every model for which a pseudotrial comparison was run.

### 2.9 Full-montage topographic analysis

To establish whether any surviving coupling was specific to the pre-specified Fz–Pz electrode pair or instead reflected a property of the signal common to the whole scalp, the key models were re-estimated across the full 33-channel montage (30 scalp electrodes plus the three electrooculogram channels). Three topographic validations were run. First, a cross-channel analysis related the early-window mean at each electrode to the later P300 at Pz, mapping where early activity forecasts the parietal component. Second, a same-channel analysis related the early-window mean at each electrode to the later-window mean at that same electrode, mapping the within-electrode early-to-late coupling. Third, a shape analysis related two representative complexity/dynamics measures (permutation entropy and Hjorth mobility) at each electrode to the P300 at Pz. For every electrode and analysis, both real-trial and Config-4 pseudotrial estimates were obtained, yielding scalp distributions of the coupling coefficient, the marginal *R*², and the autocorrelation ratio (AUR; Section 2.8).

The electrooculogram channels were retained throughout as negative controls. The logic of the comparison is that a coupling driven by stimulus-locked neural activity should weaken under pseudotrial substitution, whereas a coupling reflecting the autocorrelation structure of the signal should be preserved or strengthened; if the same-channel coupling at scalp sites behaves identically to that at the eye channels where no parietal cognitive process is plausible, then the coupling cannot be a P300-specific phenomenon. A coupling genuinely tied to the P300 generator should appear at centro-parietal sites and not elsewhere; a coupling reflecting general within-trial temporal continuity should appear at all electrodes alike. Scalp distributions were rendered as topographic maps using the MNE-Python *plot_topomap* routine with the standard 10–20 electrode layout.

### 2.10 Individual-differences analysis

For the complexity family, per-subject coupling slopes were estimated by ordinary least-squares regression of standardized P300 amplitude on the standardized early-window measure within each participant. For each measure we report the range, mean, and standard deviation of the per-subject slopes, the proportion of participants with positive slopes, and a *heterogeneity ratio* defined as the slope standard deviation divided by the absolute value of the population mean slope. A large heterogeneity ratio indicates that individual coupling slopes vary substantially in magnitude and direction relative to their average, so that the population mean is not representative of the individuals composing it. Cross-measure consistency was assessed by correlating each participant’s slopes across the three complexity measures.

### 2.11 Software and reproducibility

All analyses were conducted in Python 3. EEG processing used MNE-Python (Gramfort et al., 2013); numerical and statistical computation used *NumPy* (Harris et al., 2020) and *SciPy* (Virtanen et al., 2020); data management used pandas (McKinney, 2010); mixed-effects models were fitted in statsmodels (Seabold & Perktold, 2010); and entropy and complexity measures were computed with *antropy* (Vallat, 2023). The cross-validation dataset was accessed via *EEGDash.* All stochastic operations, including pseudotrial placement, used fixed random seeds.

Filter settings, the ICA configuration, epoch and analysis windows, the artifact-rejection threshold, the pseudotrial placement parameters and seed, and the trial-count criterion were fixed in a single configuration applied identically across datasets. The complete analysis pipeline, including the configuration files specifying every parameter reported here, will be made available in a public repository on publication.

## 3. Results

### 3.1 Overview

The paradigm produced a positive parietal response to target stimuli with the expected centro-parietal topography (grand-average waveforms at Fz and Pz, Figure 1A; scalp topography of the P300-window response, Figure 1B). With the corrected event coding the target-to-standard ratio was the expected ∼1:4 of the oddball design (213 target and 871 standard trials retained). Targets elicited a larger parietal positivity than standards, though the contrast in this sample was modest and did not reach significance (target minus standard +1.56 µV at Pz; paired *t* = 1.55, *p* = 0.13; 13 of 27 participants positive). This does not bear on the analyses reported here: every coupling model uses the within-participant standardized single-trial P300 amplitude with experimental condition included as a covariate, so the findings concern trial-to-trial covariation of the P300 rather than the target–standard difference. Figure 2 shows the rectified early-window Fz signal whose features are used as predictors. Against this backdrop, the three measure families behaved in three qualitatively different ways under pseudotrial control. Cross-channel amplitude and energy couplings strengthened under pseudotrial substitution; the same-channel coupling was large and unchanged under substitution and present at every electrode; and the complexity couplings were small at the population level, weakened under substitution, and concealed large individual-level heterogeneity. Each pattern replicated in the independent dataset. Table 4 summarizes the headline coupling estimates and their pseudotrial behavior across both datasets.

**Figure 1.**
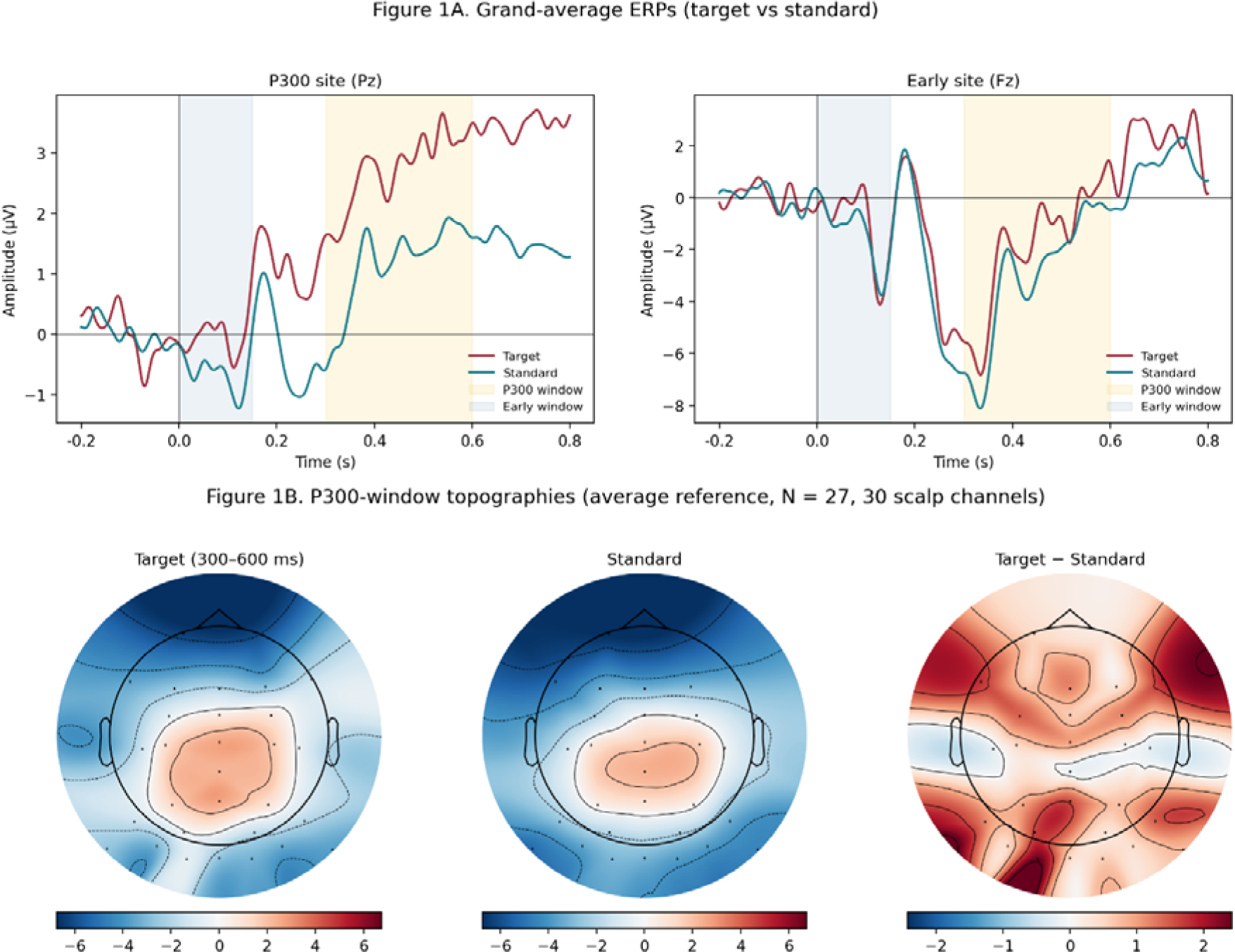
Grand-average ERPs and topographic diagnostic (primary dataset, *N* = 27). (A) Grand-average ERPs at Pz (left) and Fz (right) for target (red) and standard (blue) trials. Both the early window (0--150 ms, blue shading) and the P300 window (300--600 ms, yellow shading) are marked in both panels. Targets elicit a positive parietal deflection in the P300 window; the target--standard contrast in this sample was small and non-significant. (B) Scalp topography of activity averaged over the P300 window for target trials (left), standard trials (center), and the target-minus-standard difference (right; note the separate color scale). The difference map confirms a centroparietal positivity peaking near Pz despite the small grand-average difference.

**Figure 2.**
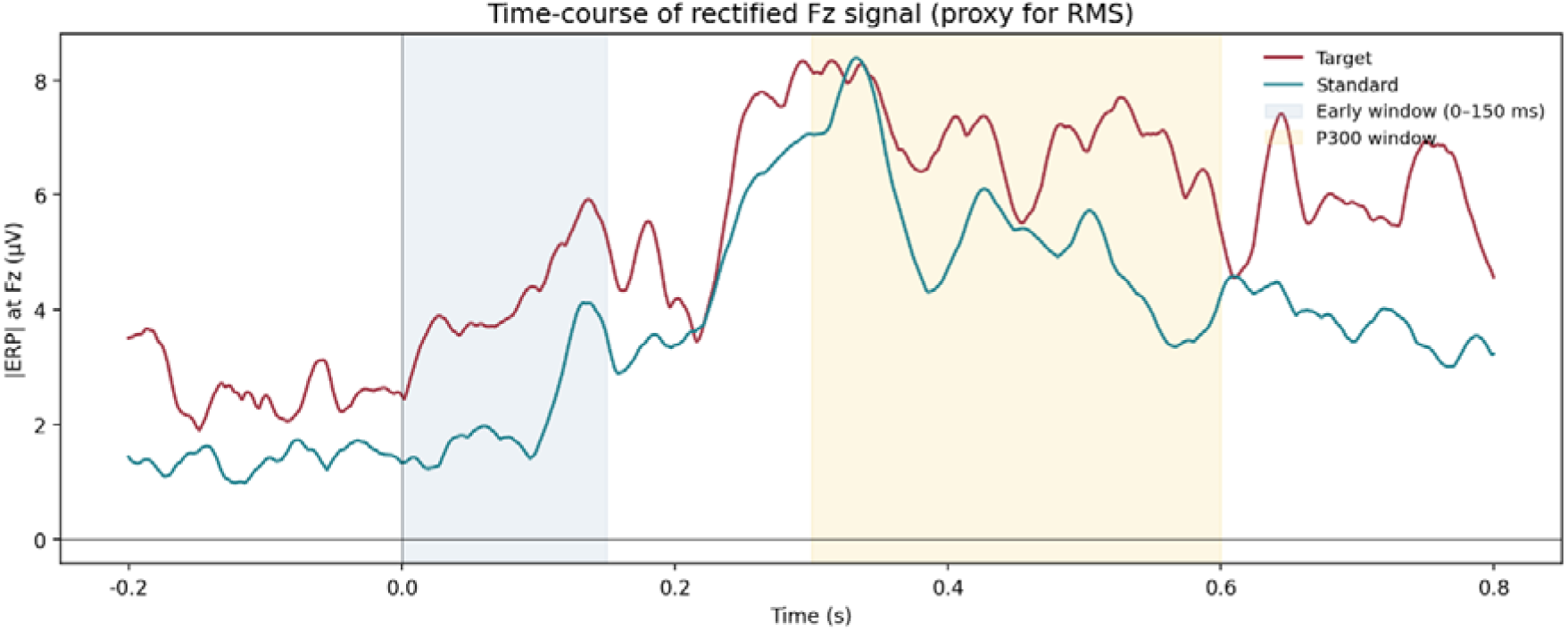
Time-course of the rectified (absolute-value) Fz signal for target (red) and standard (blue) trials, averaged within subjects and then across the 27 retained subjects, with the 0--150 ms early window shaded. The figure visualizes the early-window signal whose features (mean, RMS, entropy) are used as predictors throughout the analysis, and confirms that both conditions produce similar frontal energy in this window

**Table 4.** Headline coupling estimates and pseudotrial behavior. *R*²_real_ is the coupling in real trials; *R*²_pseudo_ is the coupling in pseudotrials; AUR is the autocorrelation ratio. Cross-validation models are estimated on *N* = 90 real-trial participants (3,130 target trials) and *N* = 83 matched-pseudotrial participants (2,270 pseudotrials).

| Model | Description | Dataset | $R^2_{\text{real}}$ | $R^2_{\text{pseudo}}$ | AUR | Interpretation |
| --- | --- | --- | --- | --- | --- | --- |
| M1 | Frontal RMS → parietal P300 (cross-channel) | Primary ($N = 27$) | 0.007 | 0.015 | 1.47 | Background autocorrelation |
| M1 | Frontal RMS → parietal P300 | ds006018 ($N = 90$) | 0.007 | 0.000 | 0.03 | Near-zero; sample-dependent |
| M9a | Same-channel temporal continuity (early Pz → late Pz) | Primary ($N = 27$) | 0.310 | 0.370 | 1.02 | Within-trial temporal continuity |
| M9a | Same-channel temporal continuity | ds006018 ($N = 90$) | 0.231 | 0.246 | 0.95 | Within-trial temporal continuity |
| M8 | Same-channel energy (RMS, early Pz → late Pz) | Primary ($N = 27$) | 0.021 | 0.066 | 1.21 | Same-channel energy (grows under substitution; weakly autocorrelation) |
| M8 | Same-channel energy | ds006018 ($N = 90$) | 0.037 | 0.002 | 0.19 | Stimulus-locked-style (collapses) |
| M13 | Permutation entropy → P300 | Primary ($N = 27$) | 0.010 | 0.002 | 0.54 | Weak, stimulus-locked at population level |
| M14 | Sample entropy → P300 | Primary ($N = 27$) | 0.003 | 0.002 | 1.11 | Weak; approximately preserved (AUR 1.11); see Section 3.5 |
| M15 | Lempel–Ziv → P300 | Primary ($N = 27$) | 0.012 | 0.006 | 0.95 | Weak; stimulus-locked (AUR $< 1$) |
| M13–M15 | Complexity → P300 | ds006018 ($N = 90$) | 0.000 | 0.000 | n/a | Population-level null (AUR undefined at $R^2 \approx 0$) |

Table 5 reports the standardized coupling coefficient (β) and marginal *R*² on real trials, and the Config-4 pseudotrial coefficient with the autocorrelation ratio (AUR; Section 2.8), for every model in the primary dataset. The pattern is consistent within each family: the amplitude and energy models carry coupling that is preserved or amplified under pseudotrial substitution (e.g., M1, AUR=2.51, M18, AUR= 2.27), the same-channel competitive model M9a shows the largest coupling of any model (*R*² = 0.310) with its coefficient unchanged under substitution (AUR=1.00), and the complexity and shape models carry weak coupling (*R*² ≤ 0.017) that does not survive as stimulus-locked.

**Table 5.** Full model results in the primary dataset. (*N* = 27, 1,084 trials; 213 target, 871 standard). β is the standardized fixed-effect coefficient for the focal predictor; *R*²_real_ is the marginal *R*² on real trials; β_pseudo_ is the coefficient on Config-4 pseudotrials; AUR is the autocorrelation ratio. The Signature column combines a coupling-magnitude label with a pseudotrial verdict. The magnitude labels describe variance explained on real trials and carry no claim of practical importance: the entire endpoint-summary family explains little of the single-trial variance, and the real-trial *R*² values fall into two clearly separated groups with nothing in between **Negligible** (*R*²_real_ ≤ 0.03; every model except the three below) and **Large** (*R*²_real_ ≈ 0.31; only the same-channel parietal-mean models M9a, M9b, M12). “Large” here means large *relative to the rest of the family*; the pseudotrial control shows even this coupling is not stimulus-locked (see Section 3.3). Within the Negligible group, the AUR and *p*-values in Sections 3.2 and 3.5 distinguish the few couplings that reach real-trial significance from those that do not. The pseudotrial verdict classifies the direction of change under substitution: **autocorrelation** (AUR > 1.20: coupling preserved or stronger on non-stimulus-locked windows), **approximately preserved** (AUR 0.80–1.20: essentially unchanged), and **stimulus-locked** (AUR < 0.80: coupling weakens when stimulus locking is removed). Where the real-trial coupling is itself near-zero the AUR is a directional qualifier only, since the ratio is unstable when its denominator is near-zero. For competitive models (M4a/M4b, M9a/M9b) *R*² reflects the full model including the covariate, so the focal-predictor β, not the overall *R*², determines the magnitude label. All models have pseudotrial estimates.

| Model | Measure / window | $\beta_{\text{real}}$ | $R^2_{\text{real}}$ | $\beta_{\text{pseudo}}$ | AUR | Signature |
| --- | --- | --- | --- | --- | --- | --- |
| *Frontal cross-channel (Fz → Pz)* |  |  |  |  |  |  |
| M1 | RMS, 0–150 ms | −0.073 | 0.007 | −0.106 | 1.47 | Weak; autocorrelation (AUR 1.47) |
| M2 | Mean, 0–150 ms | +0.106 | 0.013 | +0.223 | 2.10 | Weak; autocorrelation (AUR 2.10) |
| M3 | SD, 0–150 ms | −0.039 | 0.002 | −0.040 | 1.00 | Null; AUR ≈ 1 (preserved) |
| M4a | Mean, competitive (focal: mean) | +0.105 | 0.014 | +0.221 | 2.10 | Weak; autocorrelation (AUR 2.10; same pattern as M2) |
| M4b | SD, competitive (focal: SD) | −0.037 | 0.014 | −0.037 | 1.01 | Null focal ($R^2$ from mean covariate); AUR ≈ 1 |
| M5 | RMS, 0–200 ms | −0.033 | 0.002 | −0.113 | 3.38 | Null real; strong autocorrelation in pseudo (AUR 3.38) |
| M6 | RMS, 0–250 ms | −0.054 | 0.004 | −0.109 | 2.03 | Weak; autocorrelation (AUR 2.03) |
| M5m | Mean, 0–200 ms | +0.125 | 0.019 | +0.230 | 1.85 | Weak; autocorrelation (AUR 1.85) |
| M6m | Mean, 0–250 ms | +0.148 | 0.026 | +0.227 | 1.53 | Negligible; autocorrelation (AUR 1.53) |
| M7 | Baseline RMS (−150–0 ms) | +0.009 | 0.000 | −0.086 | 9.39 | Null real; strong autocorrelation in pseudo (AUR 9.39; baseline reference) |
| M7m | Baseline mean (−150–0 ms) | +0.037 | 0.002 | +0.062 | 1.66 | Null real; autocorrelation in pseudo (AUR 1.66; baseline reference) |
| *Parietal same-* |  |  |  |  |  |  |
| *channel (Pz → Pz)* |  |  |  |  |  |  |
| M8 | RMS, 0–150 ms | –0.139 | 0.021 | –0.168 | 1.21 | Negligible; approximately preserved (AUR 1.21; same-channel energy) |
| M9a | Mean, competitive (focal: mean) | +0.569 | 0.310 | +0.582 | 1.02 | Large (vs. family); temporal continuity, not stimulus-locked (AUR 1.02) |
| M9b | SD, competitive (focal: SD) | –0.024 | 0.310 | –0.018 | 0.74 | Negligible focal coupling (the $R^2 = 0.31$ comes from the mean covariate, not from SD) |
| M10 | Baseline RMS (–150–0 ms) | –0.036 | 0.002 | –0.141 | 3.97 | Null real; strong autocorrelation in pseudo (AUR 3.97; baseline reference) |
| M11 | RMS + baseline covariate | –0.138 | 0.021 | –0.144 | 1.05 | Negligible; approximately preserved (AUR 1.05; same-channel energy, survives baseline) |
| M12 | Mean + baseline covariate | +0.571 | 0.310 | +0.589 | 1.03 | Large (vs. family); temporal continuity, survives baseline (AUR 1.03) |
| *Complexity (Fz → Pz)* |  |  |  |  |  |  |
| M13 | Permutation entropy | +0.097 | 0.010 | +0.053 | 0.54 | Weak; stimulus-locked (AUR 0.54) |
| M14 | Sample entropy | +0.054 | 0.003 | +0.060 | 1.11 | Weak; approximately preserved (AUR 1.11) |
| M15 | Lempel–Ziv complexity | +0.125 | 0.012 | +0.119 | 0.95 | Weak; approximately preserved (AUR 0.95) |
| *Distributional shape / robust / trend / Hjorth (Fz → Pz)* |  |  |  |  |  |  |
| M16 | Skewness | +0.043 | 0.002 | +0.004 | 0.10 | Null; pseudo collapses to $\approx 0$ (AUR 0.10) |
| M17 | Kurtosis | +0.035 | 0.002 | +0.051 | 1.46 | Null; pseudo grows slightly (AUR 1.46) |
| M18 | Median | +0.102 | 0.013 | +0.232 | 2.27 | Weak; autocorrelation (AUR 2.27) |
| M19 | Median absolute deviation | –0.056 | 0.004 | –0.068 | 1.21 | Weak; approximately preserved (AUR 1.21) |
| M20 | Peak-to-peak | –0.031 | 0.001 | –0.022 | 0.69 | Null; pseudo weakens (AUR 0.69) |
| M21 | Within-window slope | +0.131 | 0.017 | +0.119 | 0.91 | Weak; approximately preserved (AUR 0.91) |
| M22 | Hjorth mobility | +0.085 | 0.008 | +0.070 | 0.82 | Weak; stimulus-locked (AUR 0.82) |

| **Model** | **Measure /**<br>**window** | $\beta_{\text{real}}$ | $R^2_{\text{real}}$ | $\beta_{\text{pseudo}}$ | **AUR** | **Signature** |
| --- | --- | --- | --- | --- | --- | --- |
| M23 | Hjorth<br>complexity | -0.035 | 0.002 | -0.044 | 1.28 | Null; pseudo grows (AUR<br>1.28) |

### 3.2 Amplitude and energy measures reflect background autocorrelation

Cross-channel amplitude and energy couplings, early frontal mean or RMS activity related to the later parietal P300, showed coupling that was equal to or stronger under pseudotrial substitution than in real trials. Because the pseudotrials contain the background temporal structure of the signal but no stimulus-locked response, a coupling that survives or grows under the substitution cannot be attributed to stimulus-locked processing; it reflects the autocorrelation structure of the continuous EEG. The apparent forecasting of the parietal component by early frontal activity is, on this evidence, a property of the background signal rather than of the neural response to the stimulus. Figure 3 shows the real-trial and pseudotrial coefficients and *R*² side by side for the flagship amplitude models (M1, M4a, M9a, M12) and entropy models (M13–M15), making the pattern visible directly: for the cross-channel models the pseudotrial bars equal or exceed the real-trial bars, whereas for the same-channel model the two are indistinguishable, and for permutation entropy (M13) and Lempel–Ziv (M15) the pseudotrial bars are smaller; sample entropy (M14) is marginal (AUR 1.11).The companion model M4b, in which SD is the focal predictor with mean held constant, showed null coupling in both real and pseudotrials (β ≈ 0, R² < 0.001), confirming that once the mean-level drift is partialled out, within-trial frontal variability carries neither genuine nor spurious coupling with P300 amplitude.

**Figure 3.**
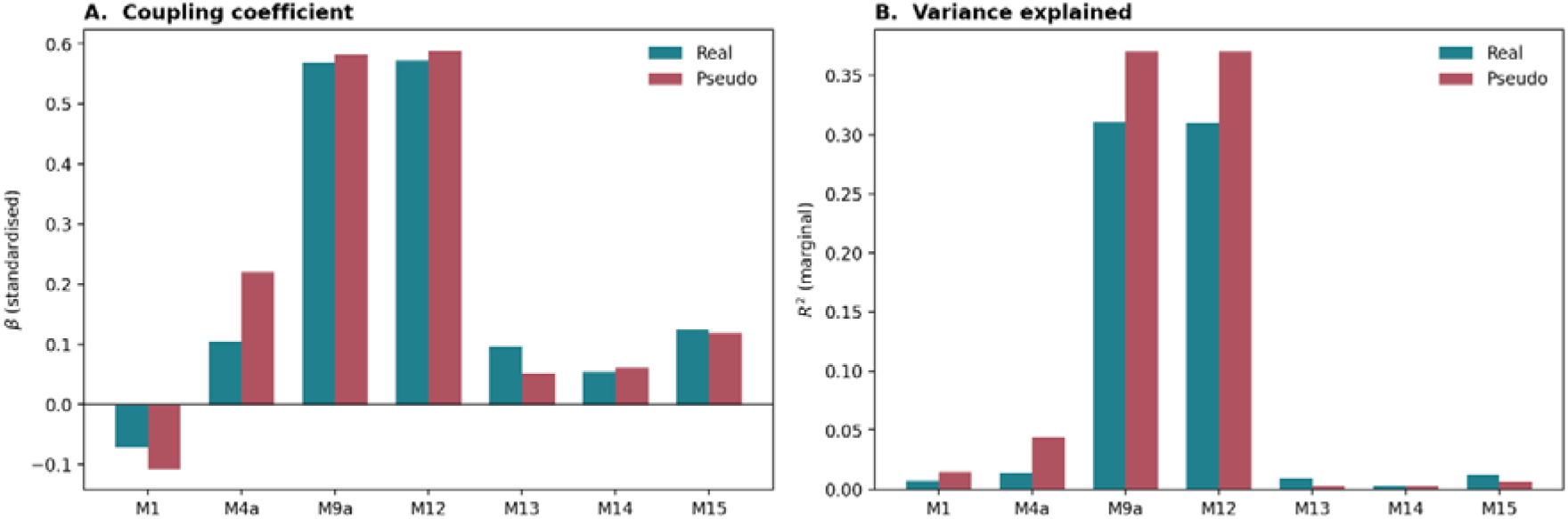
Real-trial versus pseudotrial coupling coefficients (Panel A) and marginal *R*² (Panel B) for the amplitude-family flagship models (M1, M4a, M9a, M12) and the entropy-family models (M13, M14, M15) in the primary dataset. For the cross-channel amplitude models the pseudotrial bars equal or exceed the real-trial bars (autocorrelation signature). For the same-channel model (M9a) the two bars are indistinguishable (temporal continuity). For permutation entropy (M13, AUR 0.54) and Lempel-Ziv (M15, AUR 0.95) the pseudotrial bars are smaller than the real-trial bars (stimulus-locked signature). Sample entropy (M14) is the exception: its pseudotrial coefficient is marginally larger than the real-trial coefficient (AUR 1.11), placing it in the preserved/autocorrelation category despite its near-zero coupling magnitude.

### 3.3 Same-channel coupling is general within-trial temporal continuity

The same-channel model (M9a), early-window mean amplitude at Pz related to the later P300 at the same electrode, produced the largest coupling of any model in the primary sample (β = +0.569, *R*² = 0.310). This is the kind of effect size that, taken at face value, would represent a strong single-trial predictor of the P300. Pseudotrial substitution, however, left it essentially unchanged: the coefficient was +0.582 on pseudotrials (AUR 1.02), and the marginal *R*² if anything rose slightly (0.370). By the diagnostic logic of the design, an effect whose coefficient is preserved when stimulus-locked windows are replaced by randomly placed ones is not stimulus-locked. The same conclusion held when the model included the pre-stimulus signal level as a covariate (M12: β = +0.571, *R*² = 0.310, AUR 1.03), confirming that the coupling does not depend on baseline differences between trials.

### 3.4 Topographic analysis

The full-montage analysis re-estimated the coupling models at all 33 electrodes (30 scalp electrodes plus three electrooculogram channels) on both real trials and Config-4 pseudotrials. It provides an independent, spatially resolved test of each of the three main findings: each has a predicted scalp signature, and each prediction is borne out. The three subsections below treat the three findings in turn.

#### 3.4.1 Cross-channel amplitude coupling reflects background autocorrelation

The first finding is that the apparent coupling between early activity at one electrode and the later P300 at the parietal site Pz reflects the autocorrelation of the continuous EEG rather than stimulus-locked processing. The topographic test maps this cross-channel coupling at every electrode and asks how it behaves under pseudotrial substitution. On real trials the coupling is small everywhere, with a weak focus only in the immediate vicinity of Pz (real-trial *R*² ≤ 0.094 even at centro-parietal sites and ≤ 0.03 across most of the scalp; Table 6). Decisively, it does not weaken under pseudotrial substitution but *strengthens*: at 29 of the 33 electrodes the pseudotrial coefficient exceeds the real-trial coefficient (28 of the 30 scalp sites, plus one electrooculogram channel; median AUR across the montage 1.32, rising to 1.5-1.8 at parietal and occipital sites), the Pz focus is visibly brighter on pseudotrials than on real trials, and the AUR map shows values well above 1 across most of the scalp (Figure 4). A coupling that grows when the stimulus-locked response is removed cannot be stimulus-locked; the whole-scalp AUR above one is the topographic fingerprint of background autocorrelation, matching the frontal cross-channel result of Section 3.2.

**Figure 4.**
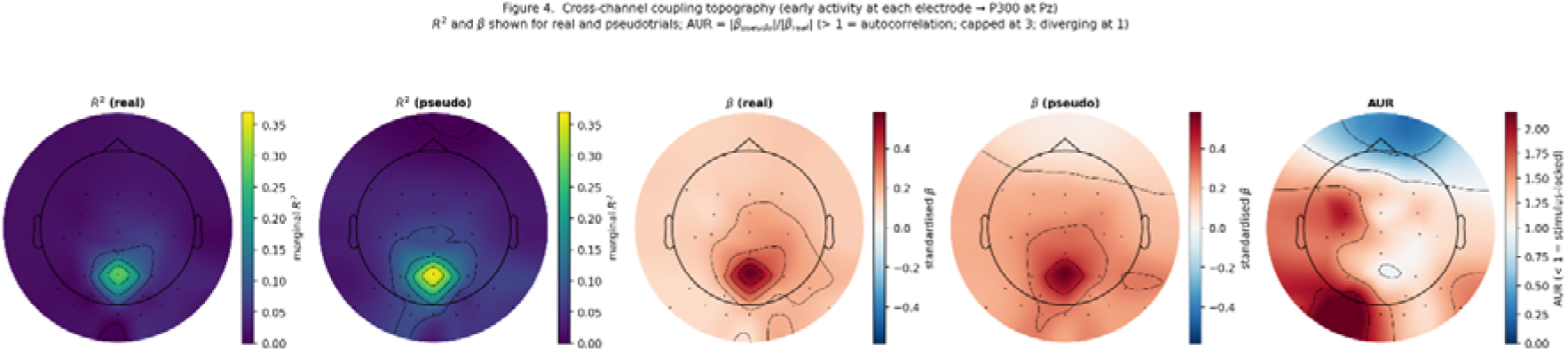
Cross-channel coupling topography (early-window mean at each electrode → P300 at Pz), primary dataset. Five panels: marginal *R*² on real trials and on pseudotrials (first two), the standardized coupling coefficient β on real trials and on pseudotrials (next two), and the AUR > 1 indicates the coupling is stronger on non-stimulus-locked windows, the autocorrelation signature; diverging scale cantered at 1). The real- and pseudotrial *R*² maps share a common color scale, as do the two β maps, so the change under substitution is read directly. The coupling is small on real trials, and strengthens under pseudotrial substitution at 29 of 33 electrodes (28 of 30 scalp sites); the two frontopolar sites (FP1: AUR 0.72, FP2: AUR 0.48) are the exception, where real-trial coupling is effectively zero. Across the remaining scalp the AUR above one confirms a background-autocorrelation rather than stimulus-locked origin.

**Table 6.**
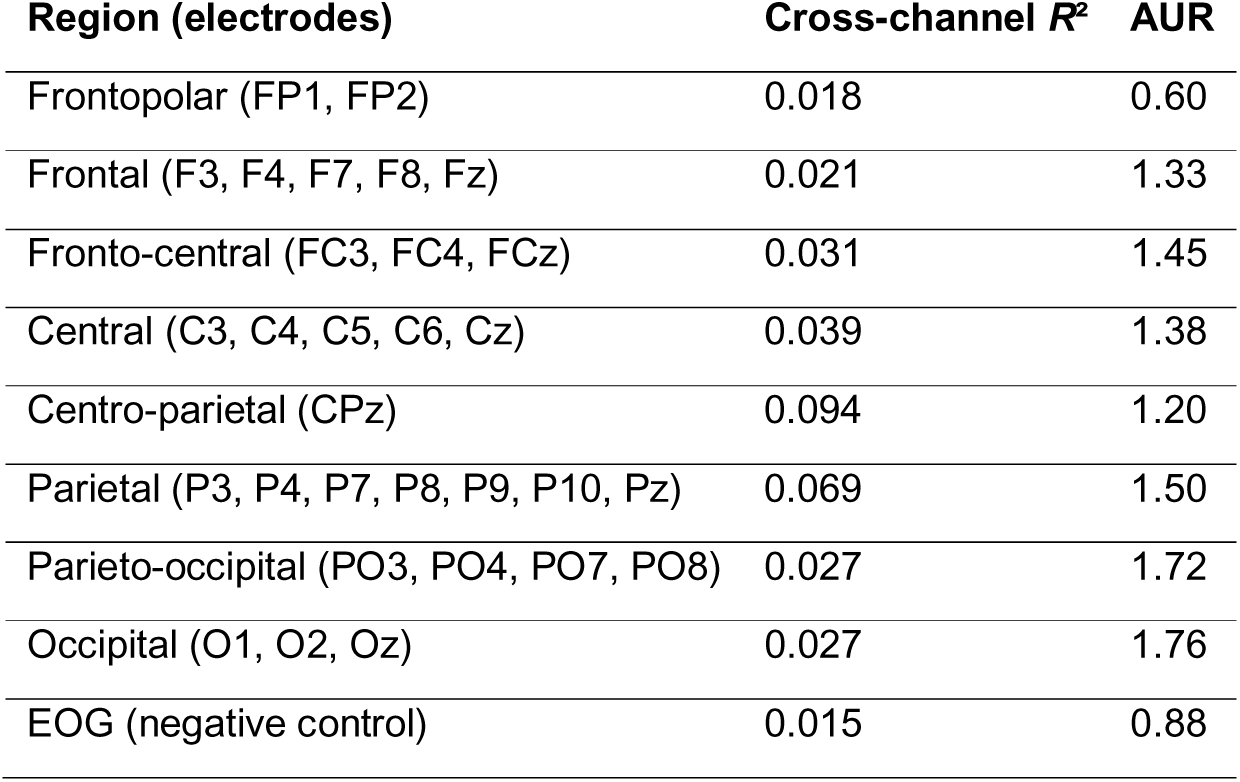
Cross-channel coupling by scalp region (primary dataset). Early-window mean at each electrode related to the later P300 at Pz. *R*² is the mean real-trial marginal *R*²; AUR is the mean autocorrelation ratio (Section 2.8). The coupling is weak everywhere and strengthens under pseudotrial substitution (AUR > 1) at 28 of 30 scalp sites; the frontopolar sites (FP1: AUR 0.72, FP2: AUR 0.48) are exceptions where coupling is effectively zero on both real and pseudotrials.

| Region (electrodes) | Cross-channel $R^2$ | AUR |
| --- | --- | --- |
| Frontopolar (FP1, FP2) | 0.018 | 0.60 |
| Frontal (F3, F4, F7, F8, Fz) | 0.021 | 1.33 |
| Fronto-central (FC3, FC4, FCz) | 0.031 | 1.45 |
| Central (C3, C4, C5, C6, Cz) | 0.039 | 1.38 |
| Centro-parietal (CPz) | 0.094 | 1.20 |
| Parietal (P3, P4, P7, P8, P9, P10, Pz) | 0.069 | 1.50 |
| Parieto-occipital (PO3, PO4, PO7, PO8) | 0.027 | 1.72 |
| Occipital (O1, O2, Oz) | 0.027 | 1.76 |
| EOG (negative control) | 0.015 | 0.88 |

#### 3.4.2 Same-channel coupling reflects within-trial temporal continuity

The second finding is that the large same-channel coupling, early-window mean at an electrode related to the later P300 at the *same* electrode (M9a, *R*² = 0.310 at Pz), is not a P300-specific parietal process but the general within-trial temporal continuity of any single-electrode signal. The topographic test re-estimates the same-channel model at every electrode: a P300-specific effect would be sharply concentrated over centro-parietal cortex, whereas signal continuity would appear everywhere. The result is unambiguous (Table 7; Figure 5). The coupling is large at every electrode, with real-trial marginal *R*² ranging from 0.175 at frontopolar sites to 0.307 at parieto-occipital sites (mean 0.233, SD 0.041). Parietal electrodes sit only marginally above the rest: the frontal midline electrode Fz, far from the P300 generator, still shows *R*² = 0.218, and the frontopolar electrodes FP1 and FP2 show *R*² ≈ 0.18. The coefficient is preserved or strengthened under pseudotrial substitution at every electrode (AUR 0.92–1.27, median 1.09), and most tellingly, the electrooculogram channels, where no parietal cognitive process is plausible, behave exactly like the scalp channels (R² ≈ 0.20, AUR 1.08–1.13). A coupling that is large everywhere, indistinguishable at the eye channels, and undiminished without stimulus locking is the temporal continuity of the ongoing signal, not a parietal P300 process.

**Figure 5.**
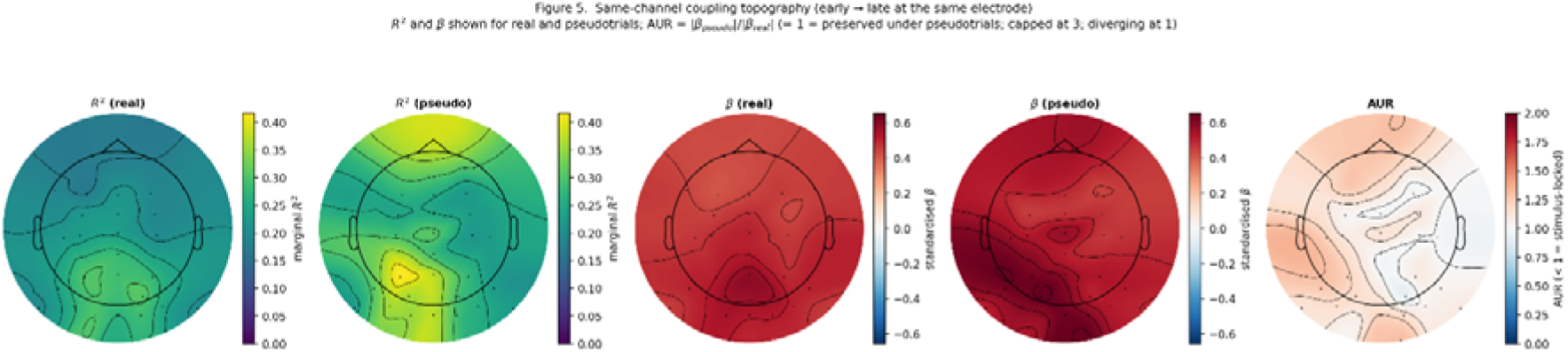
Same-channel coupling topography (early → late at the same electrode). Primary dataset. Five panels: marginal *R*² on real trials and on pseudotrials, the standardized coupling coefficient β on real trials and on pseudotrials, and the AUR; diverging scale cantered at 1). The real-and pseudotrial *R*² maps share a common color scale, as do the two β maps. The coupling is large and broadly distributed across the entire montage on real trials, preserved or strengthened at every electrode under pseudotrials (AUR 0.92-1.27), and indistinguishable between scalp and electrooculogram channels, the signature of general within-trial temporal continuity rather than a P300-specific process.

**Table 7.** Same-channel coupling by scalp region (primary dataset). Early-window mean at each electrode related to the later-window mean at that same electrode. *R*² and β are real-trial values; the AUR is the mean pseudotrial-to-real coefficient ratio (Section 2.8). The coupling is large and roughly uniform across all regions, including the electrooculogram negative-control channels.

| Region (electrodes) | Same-channel $R^2$ | Same-channel $\beta$ | AUR |
| --- | --- | --- | --- |
| Frontopolar (FP1, FP2) | 0.177 | 0.431 | 1.24 |
| Frontal (F3, F4, F7, F8, Fz) | 0.197 | 0.442 | 1.09 |
| Fronto-central (FC3, FC4, FCz) | 0.214 | 0.455 | 1.03 |
| Central (C3, C4, C5, C6, Cz) | 0.219 | 0.479 | 1.09 |
| Centro-parietal (CPz) | 0.277 | 0.516 | 0.99 |
| Parietal (P3, P4, P7, P8, P9, P10, Pz) | 0.251 | 0.504 | 1.11 |
| Parieto-occipital (PO3, PO4, PO7, PO8) | 0.282 | 0.536 | 1.07 |
| Occipital (O1, O2, Oz) | 0.286 | 0.532 | 1.15 |
| EOG (negative control) | 0.203 | 0.439 | 1.10 |

**Table 8.** Complexity coupling to the P300 across scalp regions (primary dataset). Permutation entropy and Hjorth mobility in the early frontal window, related to the later P300 at Pz. *R*²_real_ and *R*²_pseudo_ are the mean real-trial and pseudotrial marginal *R*² within each region. For the region rows the AUR_R_^2^ is the ratio of mean pseudotrial to mean real-trial *R*² (rather than the β-ratio used elsewhere), because averaging |β_pseudo_|/|β_real_| across electrodes in a region is numerically unstable when individual betas are small and near-zero (as they are for complexity measures), causing per-electrode ratios to explode; the R²-ratio aggregates stably because R² is always non-negative and gives the same directional verdict. β values exist at every electrode and are shown in Figure 6; the Fz (peak) row reports the β-based AUR used throughout the rest of the study. For both definitions, values below 1 indicate coupling that weakens under pseudotrial substitution (stimulus-locked) and values above 1 indicate autocorrelation. The frontal and central regions show stimulus-locked AUR < 1 for both measures; the high posterior AUR for permutation entropy (2.73) is an artefact of the near-zero posterior real-trial *R*² (0.0005) in the denominator, not a genuine posterior coupling, since the posterior coupling is itself negligible (see Figure 6 and the posterior real-*R*² column). Coupling is small but stimulus-locked over frontal and central sites and negligible posteriorly.

| Measure | Region / electrode | $R^2_{real}$ | $R^2_{pseudo}$ | $AUR_R^2$ | $\beta$ (real) | $p$ (real) |
| --- | --- | --- | --- | --- | --- | --- |
| Permutation entropy (M13) | Frontal (10 sites) | 0.0030 | 0.0023 | 0.76 | — | — |
|  | Central (6 sites) | 0.0025 | 0.0013 | 0.50 | — | — |
|  | Posterior (14 sites) | 0.0005 | 0.0012 | 2.73 | — | — |
| | Fz (peak electrode) | 0.0068 | 0.0022 | 0.33($AUR_{R^2}$)<br>0.59($AUR_{\beta}$) | +0.089 | 0.0008 |
| Hjorth mobility (M22) | Frontal (10 sites) | 0.0034 | 0.0022 | 0.64 | — | — |
|  | Central (6 sites) | 0.0033 | 0.0016 | 0.49 | — | — |
|  | Posterior (14 sites) | 0.0011 | 0.0011 | 1.05 | — | — |
| | Fz (peak electrode) | 0.0071 | 0.0039 | 0.55($AUR_{R^2}$)<br>0.81($AUR_{\beta}$) | +0.086 | 0.0006 |

#### 3.4.3 Complexity measures show small but reliable frontal coupling

The third finding is that the signal-complexity measures carry a small but genuine coupling to the P300, one that is anatomically specific to frontal and fronto-central electrodes and behaves as a stimulus-locked rather than an autocorrelation effect. Mapping two representative measures, permutation entropy and Hjorth mobility, at every electrode (each related to the later P300 at Pz) reveals a clear front-to-back gradient (Table 8; Figure 6). Both measures peak at the frontal midline electrode Fz, the a priori site for early frontal activity where the coupling is small in magnitude but statistically reliable (permutation entropy β = 0.089, *R*² = 0.0068, *p* = 0.0009; Hjorth mobility β = 0.086, *R*² = 0.0071, *p* = 0.0006). The effect is concentrated over frontal and fronto-central sites: across the ten frontal electrodes the coupling reached significance at six to seven of them, with mean *R*² of 0.0030 (permutation entropy) and 0.0034 (Hjorth mobility), whereas at posterior sites it was essentially absent (mean *R*² 0.0005 and 0.0011; non-significant at every posterior electrode for permutation entropy, and at all but two for Hjorth mobility, where P8 and P10 only grazed significance at *p* = 0.049 and 0.013 with *R*² ≤ 0.004). Notably, the coupling is small precisely at Pz itself (*R*² ≈ 0.000), confirming that this is not the same-channel temporal continuity of Section 3.4.2 but a distinct, frontally distributed relationship between early frontal complexity and the later parietal P300.

**Figure 6.**
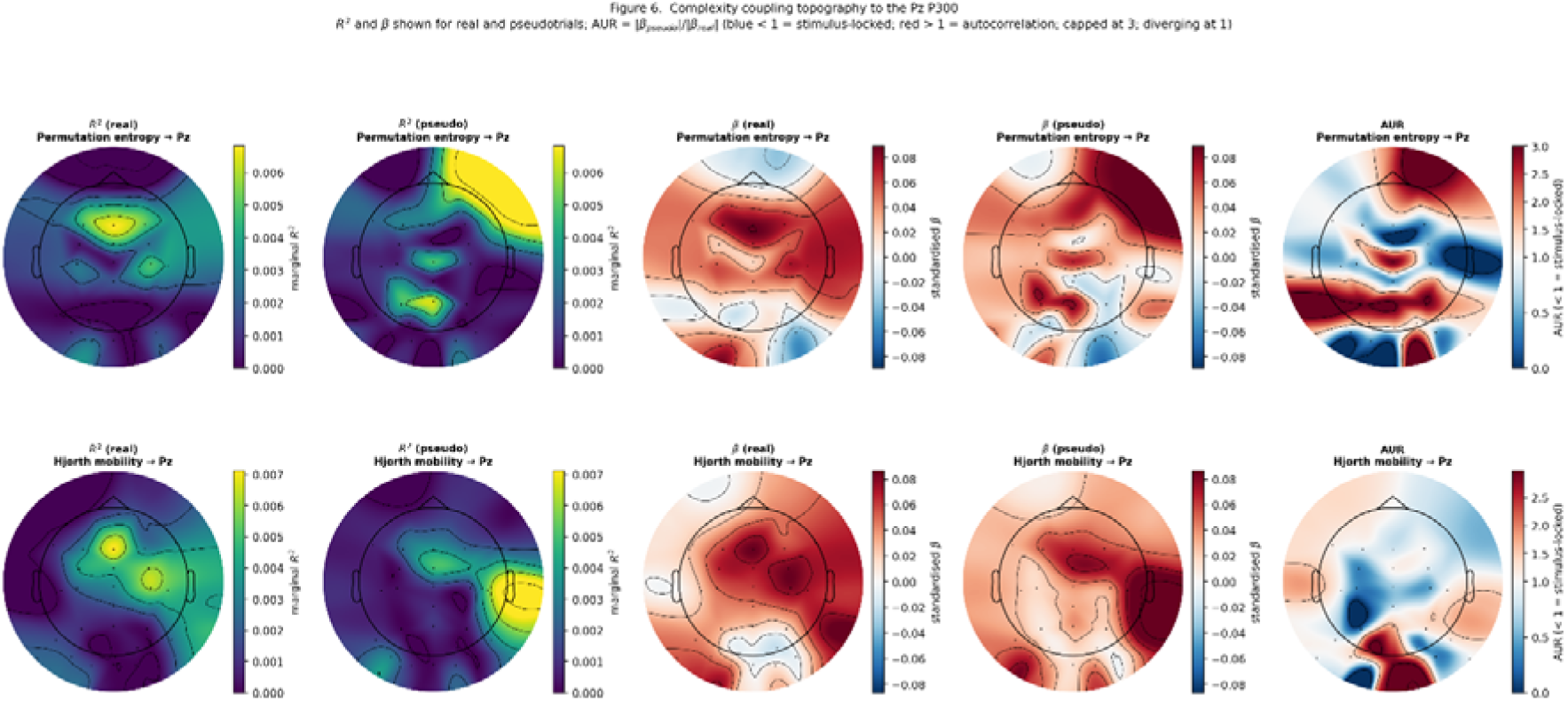
Scalp topography of complexity coupling to the Pz P300 (primary dataset). For permutation entropy (top row) and Hjorth mobility (bottom row), combining the two complexity metrics in a single figure for direct comparison. Each row shows the marginal *R*² on real trials and on pseudotrials, the standardized coupling coefficient β on real trials and on pseudotrials, and the β-based AUR; diverging scale centred at 1: blue < 1 = coupling weakens under pseudotrials = stimulus-locked; red > 1 = grows = autocorrelation; AUR capped at 3; dashed contour marks AUR = 1); within each row the real- and pseudotrial *R*² maps share a common color scale, as do the two β maps. Both measures show a small but reliable frontal/fronto-central real-*R*² focus peaking at Fz (*p* < 0.001). The AUR maps confirm the frontal coupling is stimulus-locked: Hjorth mobility shows a broad blue frontal/central region; permutation entropy shows the same at peak sites, while the red posterior region reflects near-zero real betas there. The effect is an order of magnitude smaller than the same-channel continuity of Figure 5.

Two features mark this frontal coupling as a genuine, if weak, stimulus-locked signal rather than an artefact. First, unlike the amplitude couplings of Section 3.4.1, it *weakens* under pseudotrial substitution: at Fz the permutation-entropy coefficient falls from 0.089 to 0.053 (AUR 0.59) and the Hjorth-mobility coefficient from 0.086 to 0.070 (AUR 0.81), the signature of an effect that depends on stimulus locking.

Second, it converges with the canonical frontal complexity models estimated at Fz (Section 3.5): permutation entropy (M13, *p* = 0.001) and Lempel--Ziv complexity (M15, *p* = 0.0003) both show significant, sign-consistent coupling. The topographic analysis therefore supports a modest cognitive interpretation: early frontal signal complexity carries a small amount of stimulus-locked information about the subsequent P300, consistent with the view that early-window complexity indexes the dynamical state from which stimulus processing proceeds (Section 4.2.3). The effect is real but quantitatively small, an order of magnitude below the amplitude continuity of Section 3.4.2 which is exactly why its population-level expression is modest and why the individual-differences analysis of Section 3.6 is needed to characterize it fully.

### 3.5 Complexity measures: weak population-level coupling

The signal-complexity measures behaved differently from the amplitude family. In the primary sample, the three entropy measures showed small positive population-level coupling (*R*² ≤ 0.012), and under pseudotrial substitution the coefficients tended to weaken rather than strengthen, the signature of a stimulus-locked contribution rather than of background structure. Permutation entropy (M13) showed the clearest reduction (AUR = 0.54; *R*² falling from 0.010 to 0.002) and reached significance on real trials (β = +0.097, *p* = 0.0010); Lempel–Ziv (M15) was the strongest of the three on real trials (β = +0.125, *R*² = 0.012, *p* = 0.0003) but borderline under substitution (AUR = 0.95); and sample entropy (M14) was the weakest, not reaching significance (β = +0.054, *p* = 0.09) and essentially unchanged under substitution (AUR = 1.11).

Two factors explain why SE alone shows AUR above 1. Algorithmically, at the parameters used here (*m* = 2, *r* = 0.2 × window SD), sample entropy computes conditional self-similarity over adjacent samples, a timescale at which the short-lag autocorrelation of the 1/f-filtered background EEG contributes directly to the similarity counts; ordinal-pattern and symbolic measures (permutation entropy, Lempel–Ziv) are structurally more insulated from this contribution because they encode only the rank order or discrete symbol sequence rather than distances between raw sample values. Statistically, sample entropy also has the highest individual-level sign-heterogeneity of the three measures (heterogeneity ratio 4.45 in the primary sample; 150.6 at *N* = 90, against 1.64/12.8 for permutation entropy), meaning its population mean is closest to zero by cancellation; when both the real-trial and pseudotrial population means are near-zero due to sign-heterogeneous cancellation, the AUR is unreliable and small sampling fluctuations can push the ratio above or below 1. The cross-validation confirms this: at *N* = 90 the population coupling is β = −0.009 (Table 4), indistinguishable from zero, and only the per-subject distribution reveals the genuine signal. Of the two Hjorth shape measures, mobility (M22) patterned with the entropy measures, showing a small significant real-trial coupling that weakened under substitution (β = +0.085, *R*² = 0.008, *p* = 0.004; AUR = 0.82), whereas complexity (M23) was a clear null: its coupling was negligible and non-significant on real trials (β = −0.035, *p* = 0.11) and did not change under substitution (AUR = 1.28, both coefficients near zero). In every case the effect sizes were far below any threshold for a meaningful single-trial predictor. Taken alone, the population-level analysis would have supported a modest conclusion: the complexity family carries, at most, a weak stimulus-locked signal about the P300, and not every member carries even that, sample entropy and Hjorth complexity being indistinguishable from no coupling at the population level.

### 3.6 Complexity measures: individual-differences analysis

Hjorth mobility (M22) shows a population-level stimulus-locked coupling comparable to the entropy measures (β = +0.085, *R*² = 0.008, AUR 0.82; Table 5), and its topography confirms the same frontal focus as the entropy measures (Figure 6). The individual-differences pipeline now computes per-subject Hjorth mobility and complexity slopes alongside the three entropy measures: the heterogeneity analysis saves trial-level Hjorth features for every participant in both datasets and reports their per-subject slope distributions and heterogeneity ratios on the same footing as the entropy measures (Table 9). The three entropy measures remain the primary focus of the narrative below because they index complexity in three complementary ways; the two Hjorth shape measures are reported as convergent fourth and fifth measures that show the same sign-heterogeneous structure.

**Table 9.** Per-subject coupling slopes for the entropy/complexity measures (M13–M15) and the Hjorth shape measures (M22–M23) in both datasets. The heterogeneity ratio is the slope standard deviation divided by the absolute population mean slope; larger values indicate that the population mean is less representative of individual participants. Hjorth mobility is stimulus-locked at the population level (β = +0.085, *R*² = 0.008, AUR 0.82; Table 5); Hjorth complexity shows near-zero coupling in both real and pseudotrials (β = −0.035, *R*² = 0.002, AUR 1.28; Table 5) and is accordingly classified as null rather than stimulus-locked. Both are reported here on the same per-subject basis as the entropy measures, computed by the heterogeneity pipeline (script 11), which saves trial-level Hjorth features for both datasets. All slopes use a common minimum of five clean trials per participant, so all 90 cross-validation participants contribute to every measure. The heterogeneity ratio grows with sample size for every measure (compare the primary and cross-validation rows), the defining signature of a sign-heterogeneous effect: the population mean converges toward zero as *N* increases while the spread of individual slopes does not.

| Measure | Dataset | Slope range | Mean | SD | Positive | Heterogeneity ratio |
| --- | --- | --- | --- | --- | --- | --- |
| M13 (permutation entropy) | Primary ($N = 27$) | -0.266 to +0.372 | +0.092 | 0.151 | 22/27 (81%) | 1.64 |
| M14 (sample entropy) | Primary ($N = 27$) | -0.381 to +0.459 | +0.044 | 0.197 | 16/27 (59%) | 4.45 |
| M15 (Lempel–Ziv) | Primary ($N = 27$) | -0.358 to +0.836 | +0.118 | 0.239 | 19/27 (70%) | 2.03 |
| M22 (Hjorth mobility) | Primary ($N = 27$) | -0.289 to +0.372 | +0.065 | 0.167 | 18/27 (67%) | 2.57 |
| M23 (Hjorth complexity) | Primary ($N = 27$) | -0.318 to +0.133 | -0.040 | 0.120 | 10/27 (37%) | 3.03 |
| M13 (permutation entropy) | ds006018 ($N = 90$) | -0.666 to +0.604 | +0.018 | 0.235 | 48/90 (53%) | 12.8 |
| M14 (sample entropy) | ds006018 ($N = 90$) | -0.510 to +0.585 | +0.001 | 0.186 | 44/90 (49%) | 150.6 |
| M15 (Lempel–Ziv) | ds006018 ($N = 90$) | -0.589 to +0.626 | -0.008 | 0.230 | 45/90 (50%) | 29.0 |
| M22 (Hjorth mobility) | ds006018 ($N = 90$) | -0.555 to +0.770 | +0.023 | 0.212 | 49/90 (54%) | 9.03 |
| M23 (Hjorth complexity) | ds006018 ($N = 90$) | -0.635 to +0.555 | -0.026 | 0.179 | 36/90 (40%) | 6.78 |

The per-subject analysis told a different story. For every complexity and shape measure, individual coupling slopes varied substantially in both magnitude and direction (Table 9). For permutation entropy, slopes ranged from −0.266 to +0.372 (mean +0.092, SD 0.151; 22 of 27 participants positive; heterogeneity ratio 1.6). For sample entropy, slopes ranged from −0.381 to +0.459 (mean +0.044, SD 0.197; 16 of 27 positive; ratio 4.5). For Lempel–Ziv, slopes ranged from −0.358 to +0.836 (mean +0.118, SD 0.239; 19 of 27 positive; ratio 2.0). Hjorth mobility behaved like the entropy measures (−0.289 to +0.372; mean +0.065, SD 0.167; 18 of 27 positive; ratio 2.6), and Hjorth complexity showed the same dispersion with a negative-leaning center (−0.318 to +0.133; mean −0.040, SD 0.120; 10 of 27 positive; ratio 3.0). In every case the slope standard deviation exceeded the absolute population mean, the defining feature of a population whose average is not representative of its members.

Crucially, the individual slope directions were consistent across the three measures (inter-measure correlations *r* = 0.51–0.67). Three participants showed consistent negative slopes across all three complexity measures, fourteen showed consistently positive slopes, and the remaining ten were mixed. A participant’s coupling direction was thus a stable property of that participant rather than measurement noise: the same individuals who coupled negatively on one entropy measure tended to couple negatively on the others. This within-person consistency indicates that the direction of the entropy–P300 relationship is a person-level characteristic.

The independent dataset confirmed this account directly. At the population level, all five complexity and shape measures showed coupling indistinguishable from zero in the cross-validation sample (permutation entropy β = +0.013, sample entropy β = −0.009, Lempel–Ziv β = −0.011; all *R*² ≤ 0.0002), so the modest positive population means of the primary sample did not reappear. The per-subject analysis at *N* = 90, however, revealed exactly the same sign-heterogeneous structure as in the primary data, now far more sharply resolved (Table 9; Figure 7). All measures split almost evenly between positive and negative individual slopes: permutation entropy 48 of 90 positive (53%), sample entropy 44 of 90 (49%), Lempel–Ziv 45 of 90 (50%), Hjorth mobility 49 of 90 (54%), and Hjorth complexity 36 of 90 (40%), with heterogeneity ratios of 12.8 (PE), 150.6 (SE), and 696 (LZ) for the entropy measures and 9.0 (mobility) and 6.8 (complexity) for the Hjorth measures, the slope standard deviation exceeding the near-zero population mean by one to nearly three orders of magnitude.

**Figure 7.**
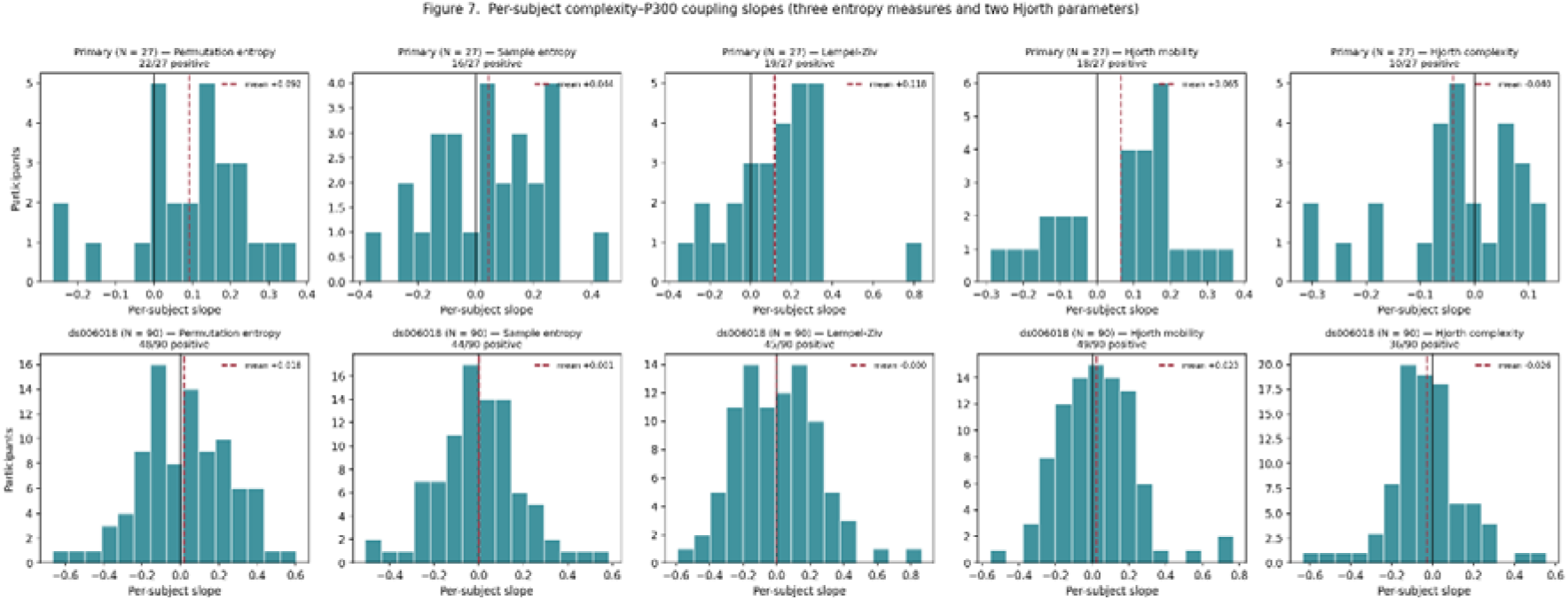
Distributions of per-subject coupling slopes for the three entropy measures (permutation entropy, sample entropy, Lempel-Ziv) and the two Hjorth parameters (mobility, complexity). In the primary dataset (top row, *N* = 27) and the cross-validation dataset (bottom row, *N* = 90). Each histogram shows one slope per participant; the dashed line marks the population mean. In the primary sample the distributions are broad and modestly positive-shifted (Hjorth complexity is the exception, with a modestly negative mean); at *N* = 90 they are cantered essentially on zero while remaining wide, the signature of a sign-heterogeneous effect whose population mean cancels despite substantial individual-level coupling. Per-subject slopes are computed directly from the trial-level data with a minimum of five clean trials per participant, so all 90 cross-validation participants are shown for every measure (Lempel--Ziv 45 of 90 positive; Hjorth mobility 49 of 90; Hjorth complexity 36 of 90).

Individual slopes again spanned more than a standardized unit (Lempel–Ziv −0.589 to +0.864). The larger sample makes the interpretation unambiguous: the modest positive means in the primary data are consistent with small-sample imbalance around a balanced distribution (drawing 19 or more positive Lempel–Ziv slopes out of 27 from a symmetric population occurs with probability 0.026), and at *N* = 90 the balance is visible directly. The individual-level coupling is real and stable within persons; its population mean is near zero because its direction varies across persons. A population-average null, where there is reason to expect sign-heterogeneous coupling, is therefore not evidence of no effect, it is evidence that the population mean is the wrong summary. The AUR panels in Figure 6 confirm that the frontal focus is genuinely stimulus-locked, not an autocorrelation artefact. The frontal sites where Figure 6 shows non-trivial *R*² (Fz, F3, F4, FC3, FC4) are predominantly blue (AUR < 1) for both measures.

### 3.7 Cross-validation of the amplitude findings in an independent dataset

The amplitude findings reproduced in the independent dataset (*N* = 90 participants, 3,130 target trials for the real-trial models; *N* = 83 participants, 2,270 pseudotrials for the Config-4 pseudotrial comparison). The same-channel temporal-continuity model M9a gave β = +0.487, *R*² = 0.231 on real trials and β = +0.462, *R*² = 0.246 on pseudotrials, an AUR of 0.95, a large coupling with the coefficient essentially preserved under pseudotrial substitution, exactly as in the primary sample. The *R*² was lower than in the primary data (0.231 vs 0.310), consistent with the differing population and recording hardware, but the defining feature, large real-trial coupling preserved under pseudotrial substitution, was fully reproduced. The same-channel energy model M8 showed the opposite, stimulus-locked-style pattern (β = +0.174 real, *R*² = 0.037; β = +0.034 pseudo, *R*² = 0.002; AUR 0.19), and the cross-channel amplitude couplings collapsed under pseudotrial substitution: the frontal RMS model M1 fell to near-zero (β = +0.077 real, +0.002 pseudo; AUR 0.03; *R*² = 0.007 on real trials), and the competitive frontal mean model M4a was weak throughout and likewise reduced (β = +0.024 real, −0.014 pseudo; AUR 0.56; *R*² = 0.0006 on real trials), consistent with these cross-channel effects carrying little stimulus-locked information in this dataset. Confirming the same-channel continuity result in an independent laboratory, population, and recording system establishes that the large M9a coupling is a structural property of the single-electrode signal rather than a feature of one dataset. (The cross-validation of the complexity findings, including the per-subject heterogeneity analysis at *N* = 90, is reported together with the primary-sample individual-differences analysis in Section 3.6; the population-level cross-validation models reported here retained 90 participants for the real-trial fits and 83 for the matched-pseudotrial fits, the small reduction reflecting participants for whom too few clean pseudotrials could be placed.)

### 3.8 Summary of results

Across two independent samples (combined *N* = 117 for the individual-differences analysis), the three measure families showed distinct and replicable signatures.

Amplitude and energy measures reflect background EEG autocorrelation, strengthening under pseudotrial substitution and varying in magnitude between samples. The large same-channel coupling reflects general within-trial temporal continuity present at every electrode, not a P300-specific parietal process, with the coefficient preserved under pseudotrial substitution in both datasets. The complexity measures carry genuine individual-level coupling whose direction is a stable person-level characteristic, producing large per-subject heterogeneity and a population mean that cancels toward zero; this sign-heterogeneous pattern was present across all five complexity and shape measures (three entropy measures plus two Hjorth parameters) in both datasets, with the within-person coupling direction correlated across the entropy measures (*r* = 0.51–0.67 in the primary sample). Not every member of the family carries even a weak population-level signal: sample entropy and Hjorth complexity were indistinguishable from no coupling at the population level, yet showed the same large per-subject dispersion, underscoring that population-level and individual-level pictures can diverge completely.

## 4. Discussion

### 4.1 Summary

We asked whether the conventional family of endpoint-summary measures captures consistent population-level stimulus-locked information about P300 amplitude in the active visual oddball once the autocorrelation structure of the EEG is controlled.

Across a primary dataset and an independent cross-validation dataset using the same paradigm, the answer is no, but the reasons differ across measure families, and one of them is more interesting than a simple null.

### 4.2 Three failure modes of endpoint-summary coupling

The results separate three distinct failure modes, each of which would, without pseudotrial control, have been read as evidence of stimulus-locked single-trial prediction. These can be grouped into three families: amplitude and energy summaries, same-channel couplings, and complexity measures.

#### 4.2.1 Failure Mode 1: Amplitude summaries and autocorrelation inflation

On the hypothesis that motivated these measures, the mean or energy of the early frontal window would index a pre-component cognitive or arousal state, the level of preparatory or attentional engagement with which the trial begins, that then shapes the magnitude of the parietal updating response, a theoretically coherent reading given that P300 amplitude scales with attentional resource allocation (Polich, 2007). Our results do not support that reading: the couplings between early activity at one site and the later P300 at another strengthened under pseudotrial substitution in the primary sample (AUR > 1) and collapsed toward zero in the larger cross-validation sample (M1 AUR 0.03 at *N* = 90), identifying them as expressions of the background autocorrelation of the continuous signal rather than of any pre-component cognitive state. The problem is not the cognitive theory but the instrument: a scalar derived by collapsing a post-stimulus window to a single value carries the 1/f autocorrelation of the ongoing signal (He, 2014; Voytek et al., 2015), and autocorrelation-inflated coupling of exactly this kind has been demonstrated formally even in EEG recorded from inanimate objects, where above-chance decoding is achieved from temporal structure with no neural signal at all (Xu et al., 2024).

The amplitude family is not internally uniform, and the pattern of differences among its members has a precise mechanistic reading (Table 10). Signed mean-level measures, mean (M2), median (M18), are the most susceptible to spurious coupling, because the mean of a short window is dominated by the lowest-frequency components of the signal, which carry the most power in a 1/f spectrum and are least attenuated by epoch-level baseline correction; spontaneous slow cortical potential (SCP) shifts within that low-frequency background are known to modulate P300 amplitude independently of the stimulus (Ergenoglu et al., 1998), producing a spurious positive coupling preserved on pseudotrials. The unsigned energy measure M1 (RMS) carries a weak negative coupling on real trials (β = −0.073) that grows more negative under pseudotrial substitution (β = −0.106; AUR = 1.47). Because RMS is dominated by unsigned high-amplitude excursions rather than by the signed DC level, it does not track the slow-drift component that inflates the signed mean-level measures; its weak negative sign reflects a broadband co-modulation in which noisier early windows accompany slightly smaller later amplitudes, and the fact that this relationship is preserved and amplified once stimulus locking is removed marks it, too, as a property of the background signal rather than of stimulus-locked processing. The standard deviation (M3) escapes both mechanisms because it is mean-centered and therefore orthogonal to slow drift (AUR ≈ 1, both coefficients near zero). The median absolute deviation (M19, AUR = 1.21) departs slightly from its parametric analogue SD (M3, AUR ≈ 1): unlike SD, which weights all deviations symmetrically around the mean, MAD is anchored to the window median and therefore retains a weak sensitivity to where that median falls within the low-frequency drift of the epoch, making it marginally more susceptible to autocorrelation inflation, though the absolute coupling remains negligible under both conditions. The competitive frontal mean model M4a behaves like the simple frontal mean M2 rather than forming an exception to it: entering the mean-centered SD as a covariate leaves the signed mean-level term essentially intact, so its coupling grows under pseudotrial substitution (β = +0.105 real to +0.221 pseudo; AUR = 2.10) to the same degree as M2 (AUR = 2.10). The slow-drift (DC/SCP) component that inflates the frontal mean lives in the signed level and is algebraically orthogonal to the mean-centered SD, so partialling SD out does not remove it, confirming that the autocorrelation inflation of M2 is carried by its DC/SCP component rather than by any feature orthogonal to SD. M4b mirrors this from the opposite direction: SD, being mean-centered, is algebraically orthogonal to slow drift, and accordingly carries null coupling under both real and pseudotrial conditions, neither spurious nor genuine. The pre-stimulus baseline models provide the cleanest demonstration of the autocorrelation mechanism, because a window measured *before* stimulus onset cannot by construction carry any stimulus-locked signal: any coupling it shows with the later P300 must be background structure. On real trials these baseline couplings are near zero (M7 frontal baseline RMS β = +0.009; M7m frontal baseline mean β = +0.037; M10 parietal baseline RMS β = −0.036), but under pseudotrial substitution they inflate sharply: the frontal baseline RMS flips sign and grows (M7 β = −0.086, AUR 9.4), while the parietal baseline RMS, already negative, inflates roughly fourfold in the same direction (M10 β = −0.141, AUR 4.0). Both trace out the same broadband-co-modulation behavior seen in the post-stimulus RMS models, expressed without any stimulus-locked component to offset it. That a pre-stimulus window can be made to “couple” with the P300 purely by the placement of the surrogate windows is the most direct possible evidence that the post-stimulus couplings of similar magnitude are autocorrelation rather than prediction. The longer-window energy variants (M5, M6) reproduce the RMS signature of M1 and the longer-window mean variants (M5m, M6m) reproduce the signed-mean signature of M2, each retaining an autocorrelation signature (AUR > 1) under pseudotrial substitution; the inflation is therefore a property of the autocorrelated background rather than of the specific 0–150 ms window length. The within-window slope (M21, AUR = 0.91) behaves like the signed mean-level family rather than the mean-centered family: slope captures the direction of low-frequency drift within the window, which is as much a property of the 1/f background structure as the DC level itself, and accordingly the coupling is largely preserved under pseudotrial substitution despite the slope’s insensitivity to absolute voltage level. The remaining distributional-shape measures, skewness (M16), kurtosis (M17), and peak-to-peak amplitude (M20), carry negligible coupling on both real and pseudotrials (all |β| ≤ 0.05, *R*² ≤ 0.002): higher-moment and range statistics of a short, heavily filtered window are dominated by noise rather than by either stimulus-locked activity or the low-frequency drift that inflates the mean-level measures, and accordingly they are simply uninformative about the P300 in either direction.

**Table 10.** Internal heterogeneity of the cross-channel amplitude family (primary dataset). Signed mean-level measures (mean, median, competitive mean) carry a slow-cortical-potential drift confound and show an autocorrelation signature (AUR > 1) with sign preserved; the unsigned RMS couples weakly and negatively and likewise grows under pseudotrial substitution (broadband co-modulation); the mean-centered SD is immune to slow drift and shows no coupling. AUR is the autocorrelation ratio (Section 2.8).

| Measure | $\beta_{\text{real}}$ | $\beta_{\text{pseudo}}$ | AUR | Mechanism |
| --- | --- | --- | --- | --- |
| Mean (M2) | +0.106 | +0.223 | 2.10 | Signed; SCP-drift driven; low-frequency dominant |
| Median (M18) | +0.102 | +0.232 | 2.27 | As mean; robust but still DC-sensitive |
| RMS (M1) | -0.073 | -0.106 | 1.47 | Unsigned; broadband co-modulation; grows under substitution |
| SD (M3) | -0.039 | -0.040 | 1.00 | Mean-centered; orthogonal to slow drifts |
| Mean, competitive (M4a) | +0.105 | +0.221 | 2.10 | Signed mean; DC/SCP drift orthogonal to SD, so partialling SD leaves it intact; grows like M2 |
| SD, competitive (M4b) | -0.037 | -0.037 | 1.01 | Mean-centered; orthogonal to slow drifts; null |
| Within-window slope (M21) | +0.131 | +0.119 | 0.91 | Slope captures the direction of low-frequency drift within the window |

#### 4.2.2 Failure Mode 2: Same-channel coupling and temporal continuity

Same-channel coupling yields the most misleading results because its effect size is exceptionally large, *R*² = 0.31 in the primary sample, 0.231 in the cross-validation sample, and large effect sizes are persuasive. Reported on its own, an early-to-late within-electrode coupling of this magnitude at Pz would look like a strong single-trial predictor of the P300. The pseudotrial control shows that it is not stimulus-locked at all: the coefficient is preserved, indeed slightly increased, when the stimulus-locked response is removed (AUR 1.00 primary, 0.95 cross-validation). As in the frontal family, the effect is carried specifically by the signed mean-level component of the window: the parietal mean (M9a, β = +0.569) drives the large coupling, whereas the parietal energy measure (M8 RMS, β = −0.139) couples in the opposite direction and the mean-centered variability (M9b SD, β = −0.024, competitive) is null. The sign dissociation is the same one seen frontally: the early-window mean tracks the slow drift that continues into the P300 window and so couples strongly and positively, while RMS, dominated by unsigned high-amplitude excursions rather than DC level, carries a weak negative relationship instead. Adjusting for the pre-stimulus baseline leaves both essentially unchanged (M12 β = +0.571, M11 β = −0.138), confirming the coupling is a property of the within-epoch signal rather than of baseline differences.

The full-montage analysis shows what the mean-level coupling is, a coupling of comparable size at every electrode (*R*² = 0.18--0.31 across the scalp, and statistically indistinguishable from scalp sites at the electrooculogram channels), reflecting the fact that the early and late portions of the same autocorrelated epoch are correlated. The baseline-shift account of P300 generation supplies a direct mechanism: if the parietal positivity is in part the visible consequence of a stimulus-driven modulation of ongoing alpha oscillations with a non-zero mean (Studenova et al., 2023), then the early and late portions of the same parietal epoch are coupled through the shared oscillatory carrier, and that carrier, alpha being distributed across the scalp, produces the same coupling at every electrode, including the eye channels. On this reading **the same-channel coupling is not a predictor of the P300 but a structural consequence of the P300’s own generation.** The lesson is methodological and general: same-channel early-to-late couplings are a trap, because the signal’s own temporal continuity guarantees a large, replicable, but entirely non-specific relationship. Effect-size magnitude is no protection; only the pseudotrial signature and the topographic ubiquity distinguish continuity from stimulus-locked coupling.

#### 4.2.3 Failure Mode 3: Complexity measures and sign-heterogeneous masking

At the population level, the complexity measures tell an unremarkable story: a weak positive coupling in the primary sample that weakens under pseudotrial substitution, and, for Lempel--Ziv complexity, a population-level coupling indistinguishable from zero in the larger cross-validation sample. A study that stopped at the population mean would conclude that permutation entropy, sample entropy, and Lempel-Ziv complexity carry essentially no single-trial information about the P300.

That conclusion would be wrong, and the per-subject analysis shows why. In both datasets the individual coupling slopes are large and broadly distributed, spanning more than a standardized unit, and they are *directionally split*: roughly half the participants couple positively and half negatively. In the larger cross-validation sample the split is almost exactly even across all five measures, permutation entropy 48 of 90, sample entropy 44 of 90, Lempel--Ziv 45 of 90, Hjorth mobility 49 of 90, and Hjorth complexity 36 of 90 positive, and the heterogeneity ratios reach 12.8, 150.6, and 29.0 for the entropy measures, the slope standard deviation exceeding the near-zero population mean by one to more than two orders of magnitude. That the heterogeneity ratio *grows* with sample size, rather than shrinking, is itself the signature of a genuine sign-heterogeneous effect: as *N* increases the population mean converges to zero while the spread of individual slopes does not. The individual-level effects are not small. They cancel at the population level because the direction of the coupling is a stable person-level characteristic that takes opposite values in different people. The within-person consistency across measures (in the primary sample, *r* = 0.51-0.67 among the three entropy measures, and as high as *r* = 0.87 between sample entropy and Hjorth mobility) confirms that this is a real, structured property of the individual, not trial-level noise: the same participants couple negatively across measures that quantify complexity in different ways.

This carries a methodological constraint that extends well beyond the present measures. A standard population-average mixed-effects model estimates a single mean coupling and will report zero for *any* measure whose coupling direction varies across individuals, regardless of how strong, real, or replicable the individual-level effect is. **The complexity-P300 relationship is a clean example: genuinely present in individuals, genuinely zero on average, and therefore invisible to the analytic tool most commonly applied to single-trial coupling questions.** Where there is reason to expect sign-heterogeneous coupling, a population-average null is not evidence of no effect; it is evidence that the wrong question has been asked of the data.

What might this person-specific coupling reflect cognitively? Signal-complexity measures of the kind used here are widely interpreted, under the neural-variability framework, as indices of the brain’s moment-to-moment dynamic range and hence of its information-processing capacity: greater signal complexity is taken to reflect a broader repertoire of accessible neural states and has been linked to more efficient and more accurate cognitive performance (McIntosh, Kovacevic, & Itier, 2008). On this view the early-window complexity of a trial is a proxy for the dynamical state the system is in as it begins to process the stimulus, and the P300 indexes the subsequent updating of the task representation (Polich, 2007); a coupling between them is therefore cognitively plausible rather than artefactual, which is consistent with our finding that the complexity coefficients weaken under pseudotrial substitution.

Two features of the complexity literature make the *direction* of that coupling a likely locus of individual differences. First, associations between single complexity measures and cognitive traits are characteristically small and vary across spatial and temporal scales, such that robust brain--behavior relations emerge only when measures are combined across individuals with cross-validation (Thiele, Richter, & Hilger, 2023). Second, where complexity does track cognition it often does so *across* individuals rather than within a fixed mapping, with high-capacity individuals distinguished by their complexity profiles during successful processing (Sheehan, Sreekumar, Inati, & Zaghloul, 2018). A single early-window complexity scalar related to a single later-component amplitude is exactly the kind of thin, scale-specific measure that this literature predicts will carry a small, person-dependent signal, **real within individuals, heterogeneous in sign across them, and therefore averaged away at the population leve**l. The present results are an instance of that general pattern, made visible by the pseudotrial and per-subject analyses rather than obscured by a population mean.

A more specific cognitive account is available for why the coupling *direction* should differ between people in a stable way. The EEG-complexity literature contains a well-replicated inversion: although greater overall brain complexity tracks greater cognitive capacity (McIntosh et al., 2008), *frontal* complexity specifically relates to performance with opposite sign for different processing modes. During convergent, focused, rule-governed thinking, high performers show *lower* frontal complexity, whereas during divergent, exploratory thinking the relationship reverses and high performers show *greater* frontal complexity (Mölle et al., 1996, 1999). Applied to the oddball task, this predicts exactly the sign heterogeneity we observe. For a participant whose habitual engagement with the task is focused-convergent, tight attentional control, reduced frontal degrees of freedom, a trial on which early frontal complexity is *lower* is a more engaged trial, yielding a larger P300 and hence a negative coupling slope; for a participant whose style is more exploratory, a higher-complexity trial is the more engaged one, yielding a positive slope. Both slopes are cognitively valid, reflecting stable individual differences in cognitive style rather than noise. That coupling direction is consistent within a person across three complexity measures that operationalize complexity in fundamentally different ways (*r* = 0.51-0.67) is the hallmark of such a trait, and is consistent with evidence that individual cortical entropy profiles are highly person-specific and stable, achieving high biometric fingerprint accuracy across recording sessions (Liu et al., 2020) and showing moderate-to-high test--retest reliability over weeks to months (Lopez et al., 2023). At *N* = 90 the population simply contains roughly equal numbers of each style, which is why the split is near-perfect and the mean is near zero.

The statistical consequence has been formalized directly: where individual effects are sign-heterogeneous, the population mean is structurally guaranteed to be near zero even when individual effects are large, stable, and replicable, so that random-slope specifications and explicit examination of the slope distribution are required to recover them (Oberauer, 2022).

Several independent lines of evidence make a balanced, two-directional split the expected outcome rather than a surprising one. At the level of the single trial, the constituent deflections of the evoked response and their cross-frequency coupling can be oppositely directed across trials and sources, so that a scalar summary of early activity need not relate to the P300 with a fixed sign (Makeig et al., 2002, 2004). The prestimulus state is itself bistable in a way that maps onto two roughly equiprobable processing regimes: the phase of ongoing oscillations at stimulus onset partitions trials into high- and low-excitability states that support opposite perceptual and cognitive outcomes (Busch et al., 2009; Dugué et al., 2011), supplying a mechanism by which the same early-window measure can precede a larger or a smaller P300 depending on the state the system happens to occupy. The cancellation is not merely an averaging accident but a known property of the coupling metrics themselves: standard phase–amplitude coupling measures, built as a distance from the uniform distribution, collapse toward zero precisely when coupling directions are opposed across the population (Tort et al., 2010). Finally, the task variables that drive the P300 are themselves balanced in a way that predicts a near-even directional split: memory and reward outcomes that modulate the P300 occur with roughly equal frequency and elicit stable but opposite individual response strategies (Sederberg et al., 2007; Yeung & Sanfey, 2004). Each of these mechanisms independently predicts that a population assembled from such individuals will contain comparable numbers coupling in each direction, exactly the ≈50/50 split observed at *N* = 90.

### 4.3 Relation to the broader single-trial literature

The three findings of this study can be read as three distinct routes by which a population-average null arises, only one of which is a genuine null and only one of which conceals recoverable cognitive content (Table 11).

**Table 11.** The three findings as three distinct origins of a population-average result. The autocorrelation ratio (AUR; Section 2.8) is the instrument that separates them when effect size alone cannot; the full-montage and per-subject analyses provide the second and third diagnostic layers.

| **Finding** | ***N*-scaling trajectory** | **AUR** | **Cognitive content** | **What the population average reports** |
| --- | --- | --- | --- | --- |
| Cross-channel amplitude (F1) | Collapses to near-zero at $N = 90$ (AUR 0.03) | $> 1$<br>(grows) | None, background autocorrelation | Correct zero |
| Same-channel continuity (F2) | Stable across $N$ | $\approx 1$<br>(preserved) | None, oscillatory continuity | Spuriously large positive |
| Complexity (F3) | Approaches exactly zero as $N$ increases | $< 1$<br>(weakens) | Real, sign-heterogeneous, trait-level | Misleading zero |

These findings urge caution in interpreting the many reports of early-window features predicting later ERP components. Where such couplings are measured within a single continuous recording and no autocorrelation control is applied, the present results show that at least three different non-stimulus-locked or non-population mechanisms can produce the appearance of prediction: background autocorrelation across channels, within-channel temporal continuity, and the accidental sign of a sign-heterogeneous individual-level effect in a small sample. The convergence with the heartbeat-evoked-potential literature, where analogous couplings proved spurious under surrogate and pseudotrial control (Steinfath et al., 2025), reinforces that the issue is not specific to one signal or paradigm but is a general hazard of single-trial coupling analysis in autocorrelated data. The same-channel result also coheres with the baseline-shift account of the P300 (Studenova et al., 2023): if the parietal positivity is in part the visible consequence of a stimulus-driven modulation of ongoing oscillations rather than a discrete additive generator, then the early and late portions of the same parietal epoch are necessarily coupled through the shared oscillatory signal, which is precisely the ubiquitous, pseudotrial-resistant, topographically non-specific coupling we observe.

### 4.4 Methodological implications

Three implications follow. **First, pseudotrial control should be a default** rather than an optional robustness check for any single-trial coupling claim in continuous EEG: the direction of change under pseudotrial substitution, not the raw effect size, is the diagnostic that separates stimulus-locked coupling from background structure.

**Second, same-channel early-to-late couplings deserve particular suspicion**, and any claim resting on one should be accompanied by a full-montage analysis demonstrating topographic specificity, because within-channel temporal continuity will otherwise masquerade as a component-specific effect. **Third, where individual-level coupling is plausibly sign-heterogeneous, analytic designs sensitive to individual differences, per-subject slopes, random-slope specifications, and explicit examination of the slope distribution, are necessary, because population-average models are structurally incapable of recovering such effects.**

### 4.5 Limitations

This study has several limitations. It examines a single paradigm, the active visual oddball, in two datasets; whether the same signatures hold in other paradigms and other components remains to be established. We did not identify the participant-level correlates of the entropy coupling direction, what distinguishes the people who couple positively from those who couple negatively is an open and interesting question that the present data cannot answer. The scope is deliberately confined to the endpoint-summary family, measures that collapse a window to a single value, and the conclusions apply to that family rather than to single-trial analysis in general. Measures that characterize the temporal *trajectory* of the early response, rather than reducing it to an endpoint, represent a categorically different approach and would require paradigms with sustained engagement to evaluate; that is a direction for intended future work.

### 4.6 Conclusion

Conventional endpoint-summary measures do not capture consistent population-level stimulus-locked P300 information in the active visual oddball once autocorrelation is controlled. Amplitude and energy measures reflect background EEG autocorrelation; the large same-channel coupling reflects general within-trial temporal continuity present at every electrode; and the complexity measures carry genuine individual-level coupling, sign-heterogeneous across persons and confirmed as such for all three entropy measures across two independent samples (combined *N* = 117 for the individual-differences analysis), whose direction therefore cancels to zero in population-average analyses. Progress on single-trial characterization of the P300 will require both instruments beyond the endpoint-summary family and analytic designs sensitive to individual differences in coupling structure.

## Data and Code Availability

Both datasets are openly available: the ERP CORE visual P3 paradigm from the ERP CORE repository (Kappenman et al., 2021) https://github.com/lucklab/ERP_CORE/tree/master/P3 and ds006018 from OpenNeuro https://openneuro.org/datasets/ds006018/versions/1.2.2 (Isbell et al., 2025), the latter distributed under a CC0 license and accessed via EEGDash. The complete analysis pipeline, all preprocessing, modelling, pseudotrial, topographic, cross-validation, and individual-differences scripts, together with the configuration files specifying every parameter reported here, is publicly available at https://github.com/erbiber/p300-entropy-metrics. Each numerical result in this paper is traceable to a specific output file through the provenance document included in the repository.

## Acknowledgments

The author thanks the ERP CORE team (Emily Kappenman, Steven Luck, and colleagues) and the DS006018 EEG dataset team (Elif Isbell, Amanda N. Peters, Dylan M. Richardson, and Nancy E. R. De León) for making their datasets publicly available, and the open-source MNE-Python community for providing robust analysis tools. This work received no external funding and reflects the author’s independent research interests at Boğaziçi University.

## Ethics Approval and Consent

Data were obtained from two publicly available EEG datasets. The ERP CORE P300 dataset (Kappenman et al., 2021) was approved by the University of California, Davis Institutional Review Board, and all participants provided written informed consent as described in the original publication. The DS006018 EEG dataset (Isbell et al. 2025) was collected in accordance with the ethical standards of the relevant institutional review board, and all participants provided informed consent as reported by the original authors. No new data were collected for the present study; accordingly, no additional ethics approval was required.

## Preprint Statement

A version of this manuscript has been posted as a preprint on bioRxiv. https://doi.org/10.64898/2025.12.17.694588

## CRediT Author Statement

Erkan Biber: Conceptualization; Methodology; Software; Formal analysis; Investigation; Data curation; Visualization; Writing: original draft; Writing: review & editing

## Supplementary Material

**Supplementary Figure S1.**
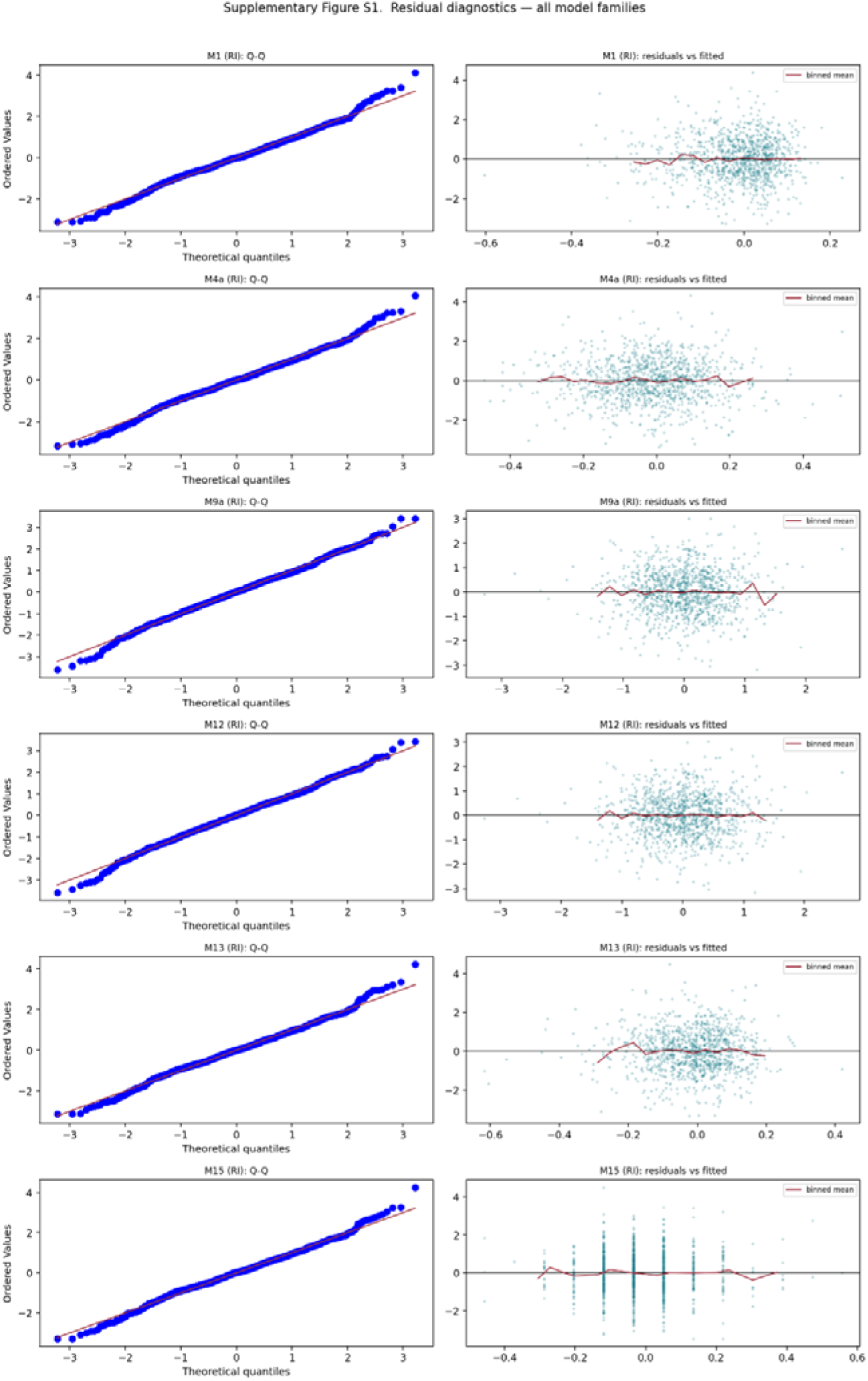
Residual diagnostics for the headline mixed-effects models across all families: amplitude (M1, M4a, M9a, M12) and entropy (M13, M15). For each model, quantile–quantile plots (left column) and residuals-versus-fitted plots (right column) confirm that model assumptions are met. The AIC comparison confirms that a random-intercept specification is preferred over a random-intercept-plus-slope for all models. Generated by script 12 (12_model_diagnostics.py).

